# Convergent generation of atypical prions in knock-in mouse models of genetic prion disease

**DOI:** 10.1101/2023.09.26.559572

**Authors:** Surabhi Mehra, Matthew E.C. Bourkas, Lech Kaczmarczyk, Erica Stuart, Hamza Arshad, Jennifer K. Griffin, Kathy L. Frost, Daniel J. Walsh, Surachai Supattapone, Stephanie A. Booth, Walker S. Jackson, Joel C. Watts

## Abstract

Most cases of human prion disease arise due to spontaneous misfolding of wild-type or mutant prion protein. Though recapitulating spontaneous prion conversion in animal models has proven challenging, transgenic mice expressing the misfolding-prone bank vole prion protein (BVPrP) recreate certain key aspects of sporadic and genetic prion disease. However, it remains unclear whether spontaneous prion generation can occur in the absence of protein over-expression and how disease-causing mutations affect prion strain properties. To address these issues, we generated knock-in mice expressing physiological levels of either wild-type or mutant BVPrP with isoleucine at codon 109. While mice expressing wild-type BVPrP remained free from neurological disease, a subset of knock-in mice expressing BVPrP with mutations that cause either fatal familial insomnia (D178N) or familial Creutzfeldt-Jakob disease (E200K) developed progressive neurological illness. Brains from spontaneously ill knock-in mice contained prion disease-specific neuropathological changes as well as atypical protease-resistant prion protein. Moreover, brain extracts from spontaneously ill D178N- or E200K-mutant BVPrP knock-in mice transmitted disease to mice expressing wild-type BVPrP. Surprisingly, the properties of the D178N- and E200K-mutant prions appeared identical both pre- and post-transmission, suggesting that both mutations guide the formation of a highly similar atypical prion strain. These findings imply that knock-in mice expressing mutant BVPrP spontaneously develop a *bona fide* prion disease and that mutations causing prion diseases may share a uniform initial mechanism of action. Therefore, these mice represent useful tools for studying the early stages of genetic prion diseases.

## Introduction

Human prion diseases such as Creutzfeldt-Jakob disease (CJD) are caused by misfolding of the cellular prion protein (PrP^C^) into PrP^Sc^, a pathological conformation that aggregates and deposits in the brain [17]. In addition to PrP^Sc^ deposition, the neuropathological hallmarks of prion disease include spongiform degeneration of the brain parenchyma as well as prominent astrocytic gliosis [71]. PrP^C^ is expressed at its highest levels in neurons and astrocytes, can be modified by the addition of up to two N-linked glycans, and is attached to the outer leaflet of the plasma membrane by a glycosylphosphatidylinositol (GPI) anchor. While PrP^C^ is predominantly α-helical, PrP^Sc^ aggregates adopt a parallel in-register β-sheet structure and are partially resistant to degradation by proteases such as proteinase K (PK) [42, 49, 51, 70]. Structural variation within PrP^Sc^ aggregates encodes distinct prion strains, which produce unique disease phenotypes [30–32, 48]. Aggregates of PrP^Sc^ can act as seeds to template the conformational conversion of PrP^C^ into additional PrP^Sc^, allowing prions to spread from cell-to-cell within the brain and from the periphery into the central nervous system.

The ability of PrP^Sc^ to self-propagate underlies the infectious nature of the prion diseases. However, most cases of prion disease in humans do not manifest due to an infectious etiology. Instead, spontaneous misfolding of PrP^C^ into PrP^Sc^ within the brain is thought to be the initiating event in sporadic prion diseases, such as sporadic CJD (sCJD). Similarly, in genetic prion disorders such as familial CJD (fCJD), fatal familial insomnia (FFI), and Gerstmann-Sträussler-Scheinker disease (GSS), mutations within the *PRNP* gene encoding PrP are believed to promote the spontaneous formation of PrP^Sc^. Different disease-causing mutations in human PrP can lead to the formation of distinct prion strains that exhibit differential susceptibility to cleavage by PK [80]. The biochemical hallmark of sCJD, fCJD, and FFI is “stereotypical” protease-resistant PrP (PrP^res^), also called PrP27-30, which is a C-terminal fragment characterized by multiple PK-resistant glycoforms with molecular weights between 19 and 30 kDa [6, 64]. In contrast, “atypical” N- and C-terminally truncated PrP^res^ fragments with molecular weights between ∼6 and 11 kDa are found in most cases of GSS [66, 77, 78].

Although ∼99% of human prion disease cases arise due to the spontaneous generation of PrP^Sc^ within the brain, little is known about the molecular mechanisms that govern this process. Inoculation of wild-type or transgenic mice with prions induces a neurodegenerative disorder that recapitulates all the biochemical and pathological hallmarks of human prion diseases [88]. However, it remains unknown whether the earliest events that occur in the sporadic and genetic prion diseases are the same as those in the infectious prion diseases, which require a pre-existing source of PrP^Sc^. Thus, there has been considerable interest in developing mouse models that exhibit spontaneous prion formation within the brain. While some success has been achieved, no existing model fully recapitulates the key features present in the human prion diseases [11, 87]. Certain lines of transgenic mice overexpressing mouse PrP (MoPrP) containing prion disease-causing pathogenic mutations develop spontaneous neurological illness but fail to generate infectious PrP^Sc^ that is highly resistant to PK digestion [10, 15, 22, 27, 34, 35, 52, 56, 79, 92]. Similarly, knock-in mouse models of fCJD, FFI, and GSS that express physiological levels of mutant MoPrP with the correct spatiotemporal expression pattern within the brain do not exhibit overt signs of neurological illness nor highly PK-resistant PrP, despite the presence of prion disease-specific neuropathological changes within the brains of certain lines [38, 39, 50]. Interestingly, transgenic mice overexpressing mutant human PrP also fail to develop neurological illness, suggesting that sequence elements within human PrP prevent spontaneous misfolding within the lifespan of a mouse [4, 5].

In recent years, bank voles (*Myodes glareolus*) have been increasingly used in prion disease research since they are susceptible to prion strains from several different species, including humans [1, 2, 21, 58, 59, 67]. Bank vole PrP (BVPrP) facilitates prion replication and formation in various *in vitro*, cellular, and animal paradigms, indicating that BVPrP is a highly permissive substrate for conversion into PrP^Sc^ [3, 13, 19, 23–26, 53, 60, 73, 85]. Moreover, transgenic mice overexpressing wild-type BVPrP containing isoleucine at polymorphic codon 109 (I109) develop a spontaneous and transmissible neurological illness characterized by prion disease-specific neuropathological changes as well as the presence of a highly PK-resistant PrP fragment in the brain [61, 84, 86]. Addition of the D178N or E200K mutations, which respectively cause FFI and fCJD, to the sequence of BVPrP(I109) hastens the onset of neurological disease in transgenic mice [84]. Collectively, these findings suggest that BVPrP(I109) is intrinsically prone to adopting misfolded, infectious conformations, potentially making it an ideal substrate for studying spontaneous prion formation under conditions of physiological protein expression.

In this study, we sought to generate improved mouse models of spontaneous prion formation by taking advantage of the unique misfolding propensity of BVPrP as well as the more translational nature of knock-in mice. Whereas knock-in mice expressing wild-type BVPrP(I109) remained healthy for their entire lifespan, knock-in mice expressing either D178N- or E200K-mutant BVPrP(I109) developed progressive signs of neurological illness, which were accompanied by the emergence of atypical PrP^res^ within the brain. Furthermore, brain extracts from spontaneously ill D178N or E200K BVPrP(I109) knock-in mice transmitted disease to knock-in mice expressing wild-type BVPrP(I109), confirming the generation of authentic prion infectivity. Surprisingly, stark differences in disease phenotype were not observed between the D178N- and E200K-mutant lines, implying that both mutations may initially act to promote the formation of a similar, perhaps identical atypical prion strain.

## Materials and Methods

### Generation and characterization of knock-in mice

Gene targeting in V6.5 embryonic stem cells was performed at the DZNE/Bonn University using CRISPR/Cas9 as described previously [40]. Plasmids containing the open reading frames (ORF) of either wild-type, D178N-mutant, or E200K-mutant BVPrP (I109 polymorphic variant) were used as a starting point [84]. Targeting constructs were generated by ligating the respective variants of the BVPrP ORF between EagI and ClaI sites of the intermediate vector pWJPrP101 [40] containing homology regions and a neomycin selection cassette removable by Flp recombinase. The Cas9 vector used for double-strand break generation in the *Prnp* gene is available from Addgene (plasmid #78621) [40]. Expansion of gene-edited embryonic stem cells and aggregation with diploid CD-1(ICR) mouse embryos was performed at The Centre for Phenogenomics (Toronto, Canada). Chimeric mice were identified by their mixed coat colors and then bred with B6(Cg)-*Tyr^c-2J^*/J mice (“B6-albino mice”; Jackson Lab #000058) to identify those that underwent germline transmission events. Chimeras were crossed with a Flp deleter strain (B6.129S4-*Gt(ROSA)26Sor^tm1(FLP1)Dym^*/RainJ; Jackson Lab #009086) to remove the selection cassette and then backcrossed with wild-type C57BL/6 mice to remove the Flp transgene. Mice that were positive for the BVPrP knock-in allele and negative for the Flp transgene were then intercrossed to create BVPrP homozygotes. All knock-in lines were maintained by crossing homozygous female with homozygous male mice.

Mice were housed at 4-5 animals per cage and were maintained on a 12 h light-12 h dark cycle, while given unlimited access to food and water. All mouse experiments were performed in accordance with the guidelines set by the Canadian Council on Animal Care under protocols (AUP #4263.17 and 6322.3) approved by the University Health Network Animal Care Committee. The presence of spontaneous neurological illness was assessed in mice 3 times per week until symptom onset. From this point, mice were assessed daily and euthanized upon progression to end-stage disease symptoms. Symptomatic profiles were determined using the standard diagnostic criteria for prion disease in mice [14]. Alternatively, mice were euthanized upon the appearance of intercurrent illness or when they reached an advanced age, typically around 600 days. Upon collection, mouse brains were divided parasagitally and then either snap frozen using dry ice and stored at −80 °C or immersed in 10% neutral buffered formalin for fixation. Mice were not perfused prior to brain collection.

### Brain homogenization and detergent extraction

Frozen mouse hemibrains were homogenized using a Minilys bead beater (PreCellys) to generate 10% (w/v) brain homogenates in Dulbecco’s phosphate-buffered saline (DPBS). For the preparation of detergent-extracted protein samples, 9 parts of 10% brain homogenate were mixed with 1 part of 10X detergent extraction buffer [5% (w/v) sodium deoxycholate and 5% (v/v) NP-40 prepared in DPBS]. The samples were incubated on ice for 20 min with vortexing every 5 min and then centrifuged at 5,000x *g* for 5 min at 4 °C. The protein concentration in the supernatant was determined using the bicinchoninic acid (BCA) assay.

### Prion transmission assays

Groups of 7-9 knock-in mice expressing wild-type BVPrP(I109) at ∼6 weeks of age were anaesthetized using isoflurane gas and then intracerebrally inoculated with 30 µL of 1% brain homogenate prepared from frozen hemibrains and diluted in PBS containing 5% (w/v) BSA. Inoculations were performed into the right cerebral hemisphere to a depth of ∼3 mm using a tuberculin syringe with an attached 27 gauge, 0.5-inch needle (BD Biosciences #305945). Inoculated mice were housed in groups of 3-4 animals in disposable cages and monitored for the development of neurological illness as described above. Once progressive signs of neurological illness were apparent, mice were euthanized, and their brains were removed and divided into hemispheres using the sagittal plane. Alternatively, the brains of asymptomatic mice were collected at the experimental endpoint (540 days post-inoculation). The left hemisphere was frozen and stored at −80 °C while the right hemisphere was fixed in 10% neutral buffered formalin and stored at 22 °C (room temperature) for neuropathological examination. Mice were not perfused prior to brain collection. All inoculation experiments utilized roughly equal numbers of male and female mice, except for one experiment involving inoculation of brain homogenate from an asymptomatic 20-month-old kiBVI^wt^ mouse, which utilized only female mice.

### Detergent insolubility assays

Detergent-extracted brain homogenates containing 50-100 μg total protein were diluted in 1X detergent extraction buffer [0.5% (w/v) sodium deoxycholate, 0.5% (v/v) NP-40 prepared in DPBS] and then ultracentrifuged at 100,000x *g* for 1 h at 4 °C using a TLA-55 rotor (Beckman). The supernatant was removed, and the pellet was resuspended in 1X Bolt LDS sample buffer (Thermo Fisher #B0007) containing 2.5% (v/v) β-mercaptoethanol. The samples were boiled at 95 °C for 10 min and then analyzed by immunoblotting.

### Enzymatic digestions

PNGase F digestions (New England Biolabs #P0704S) were performed according to the manufacturer’s recommendation. Briefly, 50 μg of detergent-extracted brain homogenate was mixed with 5 μL of Glycoprotein Denaturing Buffer (10X) and incubated at 95 °C for 10 min. The samples were allowed to cool down on ice, and then 5 μL each of 10% NP-40, 10X GlycoBuffer 2, and 0.5 μL of PNGase F were added to make a final reaction volume of 50 μL. Following incubation at 37 °C for 1-2 h, the reaction was stopped by adding LDS sample buffer (1X final concentration) and incubating the samples at 95 °C for 10 min. Samples were then analyzed by immunoblotting or ELISA.

Thermolysin (TL) was purchased from MilliporeSigma (#T7902) and dissolved in dH_2_O to generate a stock concentration of 1 mg/mL. For TL digestions, 500 μg of detergent-extracted brain homogenate was diluted into a final volume of 100 μL 1X detergent extraction buffer containing 100 μg/μL TL for a final TL:protein ratio of 1:50. The reaction mixture was incubated on a thermomixer at 37 °C for 1 h at 600 rpm, and digestions were stopped by the addition of EDTA to a final concentration of 5 mM. Sarkosyl was then added to a final concentration of 2% (v/v). This was followed by ultracentrifugation at 100,000x *g* for 1 h at 4 °C. Finally, the supernatant was gently removed, and the pellet was resuspended in 1X Bolt LDS sample buffer containing 2.5% (v/v) β-mercaptoethanol, boiled, and analyzed by immunoblotting. Alternatively, detergent-extracted brain homogenate was treated with different concentrations of TL at 37 °C for 1 h and then analyzed directly by immunoblotting without isolating the insoluble fraction. A stock solution of proteinase K (PK) at a concentration of 20 mg/mL was obtained from Thermo Scientific (#EO0491). For PK digestions, a similar protocol to that used for TL digestions was employed, except that 1 mg of detergent-extracted brain homogenate was digested with 50 μg/μL PK for a final PK:protein ratio of 1:50, and the reaction was stopped by adding PMSF to a final concentration of 2 mM.

### Immunoblotting

Proteins were separated on 10% Bolt Bis-Tris gels (Thermo Fisher Scientific) and then transferred onto 0.45 mm polyvinylidene fluoride (PVDF) membranes using Tris-Glycine transfer buffer containing 20% (v/v) methanol. Membranes were blocked in 5% (w/v) skim milk prepared in 1X Tris-buffered saline containing 0.05% (v/v) Tween-20 (TBST) overnight at 4 °C or for at least 1 h at 22 °C. The following day, membranes were incubated with primary antibody for 1 h at 22 °C. The primary antibodies that were used include anti-PrP antibodies HuM-D18 (1:5,000 dilution), HuM-P (1:10,000 dilution), HuM-D13 (1:10,000 dilution), POM1 (MilliporeSigma #MABN2285; 1:5,000 dilution), HuM-R1 (1:10,000 dilution), EP1802Y (Abcam #ab52604; 1:10,000 dilution), and SAF-32 (1:5,000 dilution); and an anti-GFAP antibody (Thermo Scientific #A-21282, 1:10,000 dilution). Blots were washed 3 times with TBST (10 min each), incubated with horseradish peroxidase (HRP)-linked secondary antibodies diluted in blocking buffer for 1 h at 22 °C, and then washed an additional 3 times with TBST. Membranes were developed using Western Lightning ECL Pro (PerkinElmer Inc.) or SuperSignal West Dura Extended Duration Substrate (Thermo Fisher Scientific) and exposure to X-ray films. For reprobing, blots were washed with TBST and then incubated in blocking buffer containing 0.05% (w/v) sodium azide overnight at 4 °C to inactivate the HRP. The next day, blots were reprobed with actin 20–33 antibody (MilliporeSigma #A5060; 1:10,000 dilution).

### Enzyme-linked immunosorbent assays (ELISAs)

Relative BVPrP expression levels in the knock-in mice were determined by ELISA. Various amounts of recombinant BVPrP(I109) [3] and detergent-extracted brain homogenates from kiBVI^wt^ mice were used as standards. Immulon 4 HBX 96-well plates (VWR #62402-959) were coated with the anti-PrP antibody HuM-D18 at 5 μg/mL in coating buffer (200 mM NaH_2_PO_4_, pH 7.5) overnight at 4 °C. The plate was blocked with 1% BSA diluted in PBS with 0.05 % Tween-20 (PBS-T) at 22 °C for > 2 h, followed by washing with PBS-T. Thereafter, the standards and test samples prepared in PBS containing 0.5 % Triton-X were added in triplicates and the plate was incubated overnight at 4 °C with shaking. Following 4 washes with PBS-T, the HRP-labeled HuM-P detection antibody was added at 1:50,000 dilution in blocking buffer and incubated at 22 °C for 2 h with shaking. The plate was thoroughly washed 5 times with PBS-T and 100 μL of TMB-Blue substrate (DAKO) was added followed by incubation in the dark for 5-10 min. The reaction was stopped by adding 100 μL of 1 M HCl to each well. Finally, the absorbance at 450 nm was read using a BMG CLARIOstar microplate reader.

### Conformational stability assays

Assays were performed using a concentration gradient of guanidine hydrochloride (GdnHCl). 20 µL of detergent-extracted brain homogenates were added to an equal volume of 2X GdnHCl stocks to create final concentrations of 1, 1.5, 2, 2.5, 3, 3.5, and 4 M GdnHCl. To generate the 0 M sample, an equal volume of DPBS was added. The samples were incubated at 22 °C for 2 h with shaking (800 rpm). The concentration of GdnHCl was then diluted to 0.4 M by adding detergent extraction buffer (1% final concentration) and DPBS. The samples were then subjected to TL digestion (100 µg/mL) as described above and ultracentrifuged at 100,000x g for 1 h at 4 °C. The supernatants were gently removed and the pellets were resuspended in 1X Bolt LDS sample buffer containing 2.5% (v/v) β-mercaptoethanol and boiled for 10 min at 95 °C. Densitometry analysis was carried out using ImageJ from three independent replicates and [GdnHCl]_50_ values (the concentration of GdnHCl at which 50% of the PrP aggregates are solubilized) were calculated using a variable slope (four parameter) dose-response model in GraphPad Prism as described previously [45].

### Neuropathology

Formalin-fixed hemibrains were processed using a Leica Pearl automated tissue processor and then embedded in paraffin. Sections (5 µm) were cut, mounted on positively charged glass slides, and then dried overnight at 37 °C. Slides were deparaffinized using xylene, rehydrated through a graded series of ethanol, and then either stained with hematoxylin and eosin (H&E) or processed for immunohistochemistry. For H&E staining, slides were stained with Hematoxylin 560 MX (Leica #3801575) for 2 min, rinsed with dH_2_O, incubated with Blue Buffer 8 (Leica #3802915) for 90 s, rinsed with dH_2_O, and then incubated with Define MX-AQ (Leica #3803595) for 30-45 s. After rinsing with dH_2_O, the slides were incubated with Eosin 515 LT (Leica #3801619) for 2 min, washed 3X with 100% ethanol (3 min each), incubated with 3 changes of xylenes (5 min each), and then coverslipped and mounted with Permount. For GFAP immunostaining, the Polink-2 Plus HRP Rabbit DAB Detection kit (GBI Labs) was used. Slides were treated with 3% (v/v) hydrogen peroxide for 10 min and then washed with dH_2_O. Antigen retrieval was performed using 10 mM sodium citrate, pH 6 for 30 min at 95 °C, and then slides were cooled and washed twice (2 min each) with TBST. The rabbit polyclonal GFAP antibody (Dako #Z0334; 1:4,000 dilution) was applied overnight at 4 °C. After rinsing with TBST, development, and counterstaining with hematoxylin, slides were dehydrated and then coverslipped and mounted using Permount Mounting Medium (ThermoFisher Scientific). For PrP immunohistochemistry, slides were treated with 98% formic acid for 10-15 min, rinsed 3 times with dH_2_O (5 min each), and then processed using the M.O.M kit (Vector Laboratories). Slides were incubated with the mouse monoclonal PrP antibody 9A2 (Wageningen Bioveterinary Research; 1:500 dilution) for 30 min at 22 °C. Slides were developed using the NovaRed system (Vector Laboratories), counterstained with hematoxylin, dehydrated, and then coverslipped and mounted using VectaMount. All slides were digitized using the Zeiss Axio Scan.Z1 slide scanner and then representative images were captured using PMA.start.

For quantification of vacuolation, snapshots of scanned slides from H&E-stained tissue were taken and then converted to 8-bit black and white images using ImageJ. Freehand regions of interest were drawn around the desired brain region, and then the threshold was set to 0-210 to reveal areas of the brain without stain. Following conversion to binary and creation of a binary mask, the Analyze Particles function in ImageJ was used to determine the percentage brain area covered by vacuolation. A size range of 8 to infinity and a circularity of 0.8 to 1.0 was used to ensure that the interiors of cerebral blood vessels were not counted as vacuoles.

For quantification of astrocytic gliosis, snapshots of scanned slides from tissue stained with a GFAP antibody were taken and converted to 8-bit black and white images using ImageJ. The H-DAB model in the IHC Toolbox plugin in ImageJ was used to remove non-stained regions. Freehand regions of interest were then drawn around the desired brain region and then the threshold was set to 0-100. The percentage area covered by GFAP staining was then measured.

### Statistical analysis

All statistical analysis was conducted using GraphPad Prism (version 10.0.0) with a significance threshold of P < 0.05. No tests for data normality were performed. Survival curves were compared by the log-rank (Mantel-Cox) test and sex-specific differences were analyzed by the Mann-Whitney test. ELISA measurements were compared using Welch ANOVA followed by Dunnett’s T3 multiple comparisons test. Detergent-insoluble PrP levels were compared using one-way ANOVA followed by Tukey’s multiple comparisons test. Neuropathological indicators were compared using a Kruskal-Wallis test followed by Dunn’s multiple comparisons test.

## Results

### Generation of BVPrP knock-in mice

To generate mice expressing physiological levels of wild-type or mutant BVPrP, we replaced the open reading frame encoding MoPrP, which is located entirely within the third exon of the *Prnp* gene, with that of BVPrP using a CRISPR/Cas9-based gene editing approach (**Fig. 1a**) [40]. Following removal of N- and C-terminal signal sequences, the amino acid sequence of mature BVPrP differs from MoPrP at 8 positions. To promote spontaneous prion formation in the brain, we used the I109 polymorphic variant of BVPrP since transgenic mice over-expressing wild-type (wt) BVPrP(I109) develop spontaneous disease much more rapidly than mice over-expressing BVPrP with methionine at codon 109 (M109) [41, 86]. We have previously found that addition of the D178N and E200K mutations to BVPrP(I109) hastens the onset of spontaneous neurological illness in transgenic mice [84]. Thus, we generated knock-in mice expressing either wt, D178N-mutant, or E200K-mutant BVPrP(I109), which will be referred to as kiBVI^wt^, kiBVI^D178N^, and kiBVI^E200K^ mice, respectively. All mice were homozygous for the knock-in alleles and thus only express wt or mutant BVPrP(I109). The E200K mutation causes fCJD when paired with either methionine (M) or valine (V) at polymorphic codon 129 in human PrP [28, 33]. In contrasting, the D178N mutation causes either FFI if it occurs *in cis* to M129 or fCJD if it occurs *in cis* to V129 [29]. Since BVPrP contains methionine at codon 129, the kiBVI^D178N^ mice are more reflective of FFI than fCJD.

**Figure 1.**
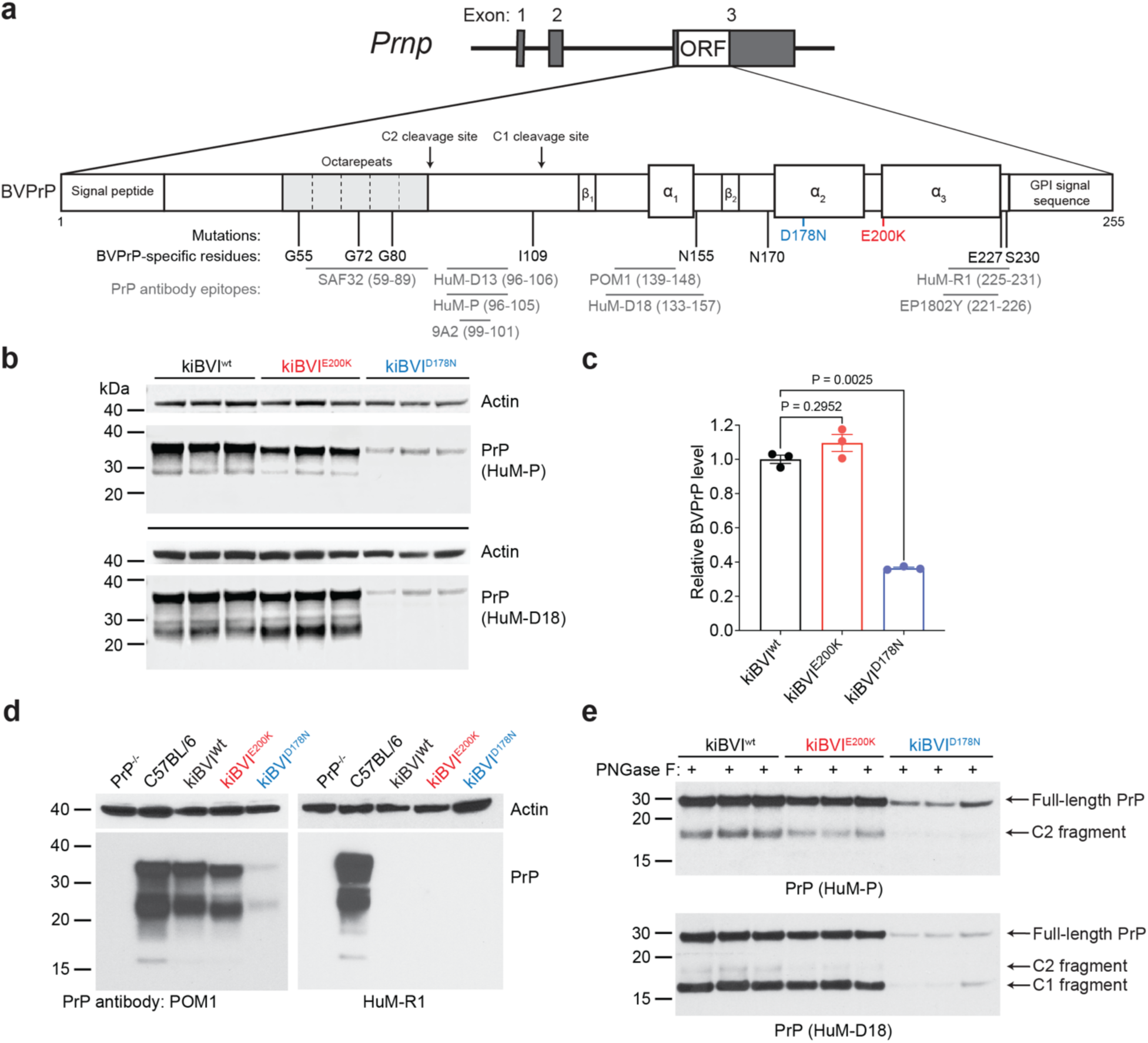
Generation and characterization of knock-in mice expressing wild-type or mutant bank vole PrP. **a**) Schematic of the gene-targeted alleles in knock-in mice expressing either wild-type (kiBVI^wt^), E200K-mutant (kiBVI^E200K^), or D178N-mutant (kiBVI^D178N^) BVPrP(I109). The eight amino acid residue differences between the mature forms of bank vole and mouse PrP are shown, as are the approximate epitopes for the anti-PrP antibodies used in this study. **b**) Immunoblots for PrP in brain extracts from 3 mice each for the indicated knock-in lines. BVPrP was detected using the antibodies HuM-P and HuM-D18, and both blots were reprobed with an anti-actin antibody. **c**) ELISA-based quantification of relative BVPrP levels (mean ± SEM) in brain extracts from knock-in mice (n = 3 per line). Statistical significance was assessed using Welch ANOVA followed by Dunnett’s T3 multiple comparisons test. **d**) Immunoblots for PrP in brain extracts from the indicated mouse lines probed with antibodies that recognize both mouse and bank vole PrP (POM1) or only mouse PrP (HuM-R1). Both blots were reprobed with an antibody against actin. **e**) Immunoblots for PrP in PNGase F-treated brain extracts from 3 mice each for the indicated knock-in lines. BVPrP was detected using the antibodies HuM-P and HuM-D18. Full-length BVPrP as well as the C1 and C2 endoproteolytic products are indicated.

Levels of BVPrP(I109) expression in the brain differed among the three lines, with relative levels dependent on the anti-PrP antibody used for detection by immunoblotting (**Fig. 1b, S1a**). PrP levels in kiBVI^E200K^ mice were similar to or slightly lower than levels in kiBVI^wt^ mice, whereas levels in kiBVI^D178N^ mice were considerably lower. Quantification of relative BVPrP(I109) expression levels in the brain by ELISA revealed that PrP levels were ∼50-60% lower in kiBVI^D178N^ mice than in kiBVI^wt^ mice but similar between kiBVI^E200K^ and kiBVI^wt^ mice (**Fig. 1c**). This is consistent with results from MoPrP-based knock-in mice and our previous finding that D178N-mutant BVPrP(I109) has a shorter half-life than either wt or E200K-mutant BVPrP(I109), resulting in lower steady-state levels [39, 84]. Brain BVPrP(I109) levels in kiBVI^wt^ mice were comparable to MoPrP levels in the brains of wild-type C57BL/6 mice, and PrP was not detected in any of the knock-in lines when using an antibody that recognizes MoPrP but not BVPrP (**Fig. 1d**). Following removal of N-linked glycans, the endoproteolytic processing of BVPrP(I109) to produce C1 and C2 fragments was observed in all 3 lines (**Fig. 1e, S1b**). This indicates that the synthesis and processing of BVPrP(I109) in the knock-in lines occurs similarly to MoPrP in wild-type mice.

### Mutant BVPrP knock-in mice develop spontaneous disease

Groups of kiBVI^wt^, kiBVI^D178N^, and kiBVI^E200K^ mice were monitored longitudinally for the development of signs of spontaneous neurological illness consistent with prion disease. Mice were analyzed up to 20 months of age since non-specific causes of morbidity or mortality in aged mice appear more frequently after this point. Whereas all kiBVI^wt^ mice remained free of neurological illness for the duration of the experiment, a subset of kiBVI^D178N^ and kiBVI^E200K^ mice developed progressive signs of neurological disease beginning around 400 days of age (**Fig. 2a**, **Table 1**). In spontaneously sick kiBVI^E200K^ and kiBVI^D178N^ mice, the most common signs of illness were prominent kyphosis and weight loss with the variable presence of dehydration, limb clasping, and gait issues. Some mice also exhibited a severe dermatitis phenotype characterized by neurotic scratching/over-grooming of the head region, and this was more common in the kiBVI^D178N^ line. Approximately 60% of kiBVI^D178N^ and kiBVI^E200K^ mice developed spontaneous disease by 20 months of age, and there was no difference in kinetics of disease onset between the two mutant BVPrP-expressing lines. For the kiBVI^D178N^ and kiBVI^E200K^ mice that developed spontaneous neurological illness, there was no difference in the age of disease onset between male and female animals (**Fig. 2b**).

**Figure 2.**
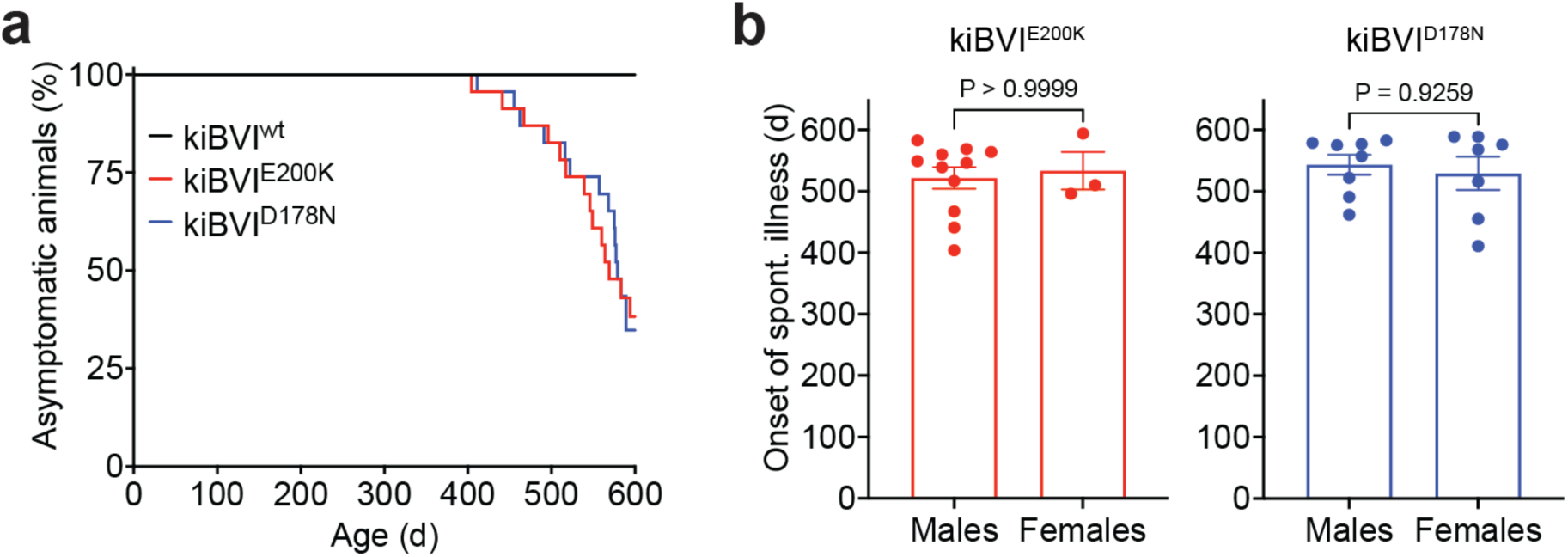
Knock-in mice expressing mutant bank vole PrP develop spontaneous neurological illness. **a**) Kaplan-Meier curves for the development of signs of neurological illness in kiBVI^wt^ (black, n = 22), kiBVI^E200K^ (red, n = 23), and kiBVI^D178N^ (blue, n = 23) mice. The onset of neurological illness in kiBVI^E200K^ and kiBVI^D178N^ mice was significantly different from kiBVI^wt^ mice (*P* < 0.0001 as assessed by the Log-rank test), whereas there was no difference in disease onset between the kiBVI^E200K^ and kiBVI^D178N^ lines (*P* = 0.97). **b**) Analysis of sex-specific effects on the age of onset of spontaneous neurological illness (mean ± SEM) in kiBVI^E200K^ (left graph; n = 11 for males, n = 3 for females) and kiBVI^D178N^ (right graph; n = 8 for males, n = 7 for females). Statistical significance was assessed using unpaired, two-tailed Mann-Whitney tests.

**Table 1.** Spontaneous neurological illness in knock-in mice expressing mutant BVPrP.

| Line | Mean age of disease onset ± SEM (d) | Signs of neurological illness (n/n<sub>0</sub>) | Mice showing TL-resistant PrP (n/n<sub>0</sub>) | Mice showing PK-resistant PrP (n/n<sub>0</sub>) |
| --- | --- | --- | --- | --- |
| kiBVI<sup>wt</sup> | > 598-603 | 0/22 | 0/21 | 0/2 |
| kiBVI<sup>E200K</sup> | 524 ± 15 | 14/23 | 14/22 | 2/6 |
| kiBVI<sup>D178N</sup> | 537 ± 15 | 15/23 | 22/22 | 7/11 |

n, number of mice; n<sub>0</sub>, number of mice examined

### Progressive accumulation of misfolded PrP species in mutant BVPrP knock-in mice

We have previously shown that spontaneously ill transgenic mice overexpressing wild-type or mutant BVPrP(I109) contain atypical PrP^res^ species in their brains [84]. Thus, we analyzed brain extracts from the knock-in mice for the presence of detergent-insoluble and protease-resistant PrP (**Fig. 3a**). Initially, we used the protease thermolysin (TL), which has been shown to effectively discriminate between normal and aggregated forms of proteins in prion disease and other protein misfolding disorders [20, 44, 52, 62]. In brains from healthy 3-month-old knock-in mice, full-length PrP species were fully degraded by TL concentrations of higher than 2 µg/mL, whereas complete digestion of the C1 endoproteolytic fragment in the kiBVI^wt^ and kiBVI^E200K^ lines required TL concentrations of at least 20 µg/mL (**Fig. S2**). Interestingly, the C1 fragment in kiBVI^D178N^ mice was much more sensitive to TL digestion, potentially suggesting that the D178N mutation destabilizes the C-terminal domain of BVPrP(I109). To ensure complete digestion of all properly folded PrP species, we used TL at a concentration of 100 µg/mL in all subsequent protease digestion experiments.

**Figure 3.**
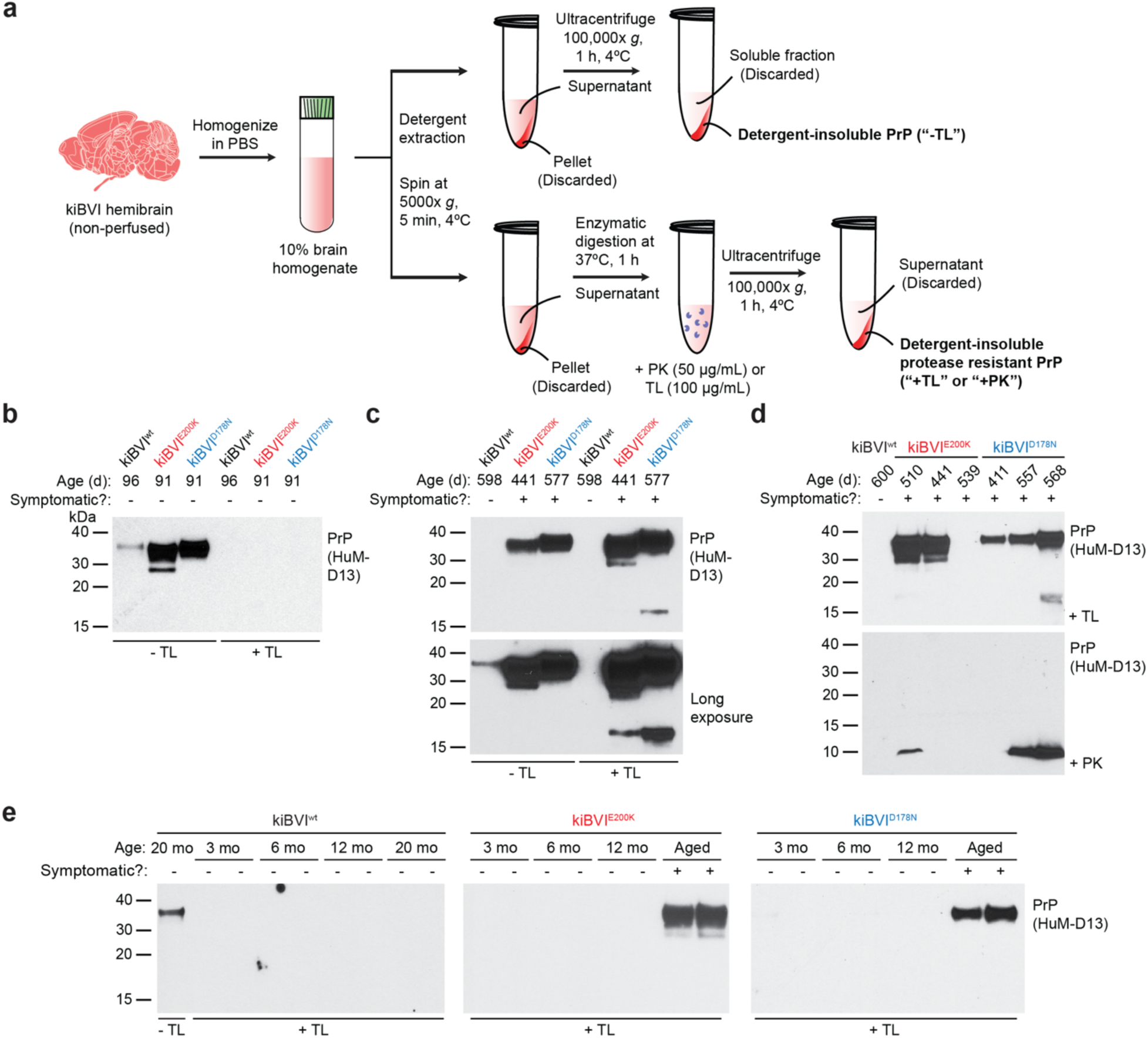
Protease-resistant PrP in brains from spontaneously ill knock-in mice expressing mutant bank vole PrP. **a**) Schematic of protocols used for the analysis of detergent-insoluble and protease-resistant PrP species in the brains of knock-in mice. **b**) Immunoblots for detergent-insoluble PrP species, with (+) or without (-) digestion with 100 µg/mL thermolysin (TL), in brain homogenates from 3-month-old kiBVI^wt^, kiBVI^E200K^, and kiBVI^D178N^ mice. **c**) Immunoblots for detergent-insoluble PrP species, with or without TL digestion, in brain homogenates from asymptomatic 20-month-old kiBVI^wt^ mice and spontaneously ill kiBVI^E200K^ and kiBVI^D178N^ mice. The bottom panel displays a longer exposure of the blot shown in the top panel. **d**) Immunoblots for detergent-insoluble PrP species in brain extracts from aged asymptomatic kiBVI^wt^ mice as well as spontaneously ill kiBVI^E200K^ and kiBVI^D178N^ mice following digestion with either 100 µg/mL TL (top panel) or 50 µg/mL proteinase K (PK, bottom panel). **e**) Immunoblots for detergent-insoluble, TL-resistant PrP species in brain extracts from kiBVI^wt^ (left), kiBVI^E200K^ (middle), and kiBVI^D178N^ (right) mice at the indicated ages. Two independent mice per age are shown. For the kiBVI^wt^ line, a sample without TL digestion is also shown. In all panels, PrP was detected using the antibody HuM-D13.

Despite expressing similar or lower levels of BVPrP, levels of detergent-insoluble PrP species were substantially elevated in 3-month-old kiBVI^E200K^ and kiBVI^D178N^ mice compared to kiBVI^wt^ mice (**Fig. 3b**). At this age, all detergent-insoluble PrP species were sensitive to TL digestion. Compared to 20-month-old kiBVI^wt^ mice, increased amounts of detergent-insoluble PrP species were also present in spontaneously ill kiBVI^E200K^ and kiBVI^D178N^ mice, and these species were resistant to TL digestion (**Fig. 3c**). TL-resistant PrP was not detected in any of the brains from 20-month-old kiBVI^wt^ mice (**Fig. 3c, d**, **Table 1**). In contrast, 100% of the brains from kiBVI^D178N^ mice that developed spontaneous neurological illness displayed TL-resistant PrP (**Fig. 3d**, **Table 1**). Moreover, all brains from asymptomatic kiBVI^D178N^ mice collected at 20 months of age also contained TL-resistant PrP (**Fig. S3**). The presence of TL-resistant PrP was more variable in the kiBVI^E200K^ line with only 64% of brains examined exhibiting clear evidence of TL-resistant species (**Table 1**). Not all kiBVI^E200K^ mice that developed spontaneous neurological illness possessed TL-resistant PrP (**Fig. 3d**), possibly due to misdiagnosis, and TL-resistant PrP was variably present in the brains from asymptomatic kiBVI^E200K^ mice collected at 20 months of age (**Fig. S3**). The main TL-resistant PrP species appeared to be similar in molecular weight to undigested PrP (**Fig. 3c, d**), and the slight difference in appearance between the kiBVI^E200K^ and kiBVI^D178N^ lines could reflect differences in N-glycosylation efficiency, which was also observed in PrP^C^ species from young mice (**Fig. 1b**), or mutation-specific differences in protein charge. A smaller and less abundant ∼17 kDa TL-resistant fragment was also observed in some of the spontaneously ill kiBVI^E200K^ and kiBVI^D178N^ mice (**Fig. 3c**).

For all three knock-in lines, total PrP levels remained largely unchanged between young and old mice (**Fig. S4a**), whereas levels of detergent-insoluble PrP were modestly increased in spontaneously ill kiBVI^E200K^ and kiBVI^D178N^ mice compared to younger mice (**Fig. S4b, c**). TL-resistant PrP was absent in kiBVI^E200K^ and kiBVI^D178N^ mice up to 12 months of age, suggesting that it may arise relatively late in the disease course (**Fig. 3e**). The brains of mice with high levels of TL-resistant PrP were also examined for the presence of PK-resistant PrP species. A PK concentration of 50 µg/mL was utilized, as this results in a rigorous digestion of PrP^C^. A subset of mice examined exhibited a PK-resistant PrP fragment with a molecular weight of ∼10 kDa (**Fig. 3d**). As with TL digestion, PK-resistant PrP species were more commonly found in brains from the kiBVI^D178N^ line than the kiBVI^E200K^ line (**Table 1**). Collectively, these results suggest that PrP^res^ progressively accumulate with age in the brains of kiBVI^E200K^ and kiBVI^D178N^ mice, but not kiBVI^wt^ mice.

### Prion disease-specific neuropathological changes in mutant BVPrP knock-in mice

Brains from spontaneously ill kiBVI^E200K^ and kiBVI^D178N^ mice exhibited prominent grey matter vacuolation in the hippocampus, cortex, and thalamus, whereas no vacuolation was observed in the brains of asymptomatic 20-month-old kiBVI^wt^ mice (**Fig. 4a, b**). Age-related white matter vacuolation was present in the brains of all older mice, regardless of symptomatic status or genotype (**Fig. S5**). There was no difference in the extent or localization of vacuolation between symptomatic kiBVI^E200K^ and kiBVI^D178N^ mice. The brains of symptomatic kiBVI^E200K^ and kiBVI^D178N^ mice also exhibited increased astrocytic gliosis compared to aged asymptomatic kiBVI^wt^ mice in the brain regions with vacuolation, as evidenced by GFAP immunolabeling (**Fig. 4c, d**). Although the increase in GFAP staining was less prominent for the kiBVI^D178N^ line, there were no statistically significant differences in the extent of GFAP immunolabeling between the kiBVI^E200K^ and kiBVI^D178N^ mice (**Fig. 4d**). Increased GFAP levels in aged symptomatic kiBVI^E200K^ and kiBVI^D178N^ mice compared to aged asymptomatic kiBVI^wt^ mice and young asymptomatic kiBVI^E200K^ and kiBVI^D178N^ mice were also observed by immunoblotting (**Fig. 4e, f**). Endoproteolytic cleavage products of GFAP similar to those found in the brains of prion-infected mice and believed to be generated by calpain cleavage were also observed in brain homogenates from symptomatic kiBVI^E200K^ and kiBVI^D178N^ mice [91].

**Figure 4.**
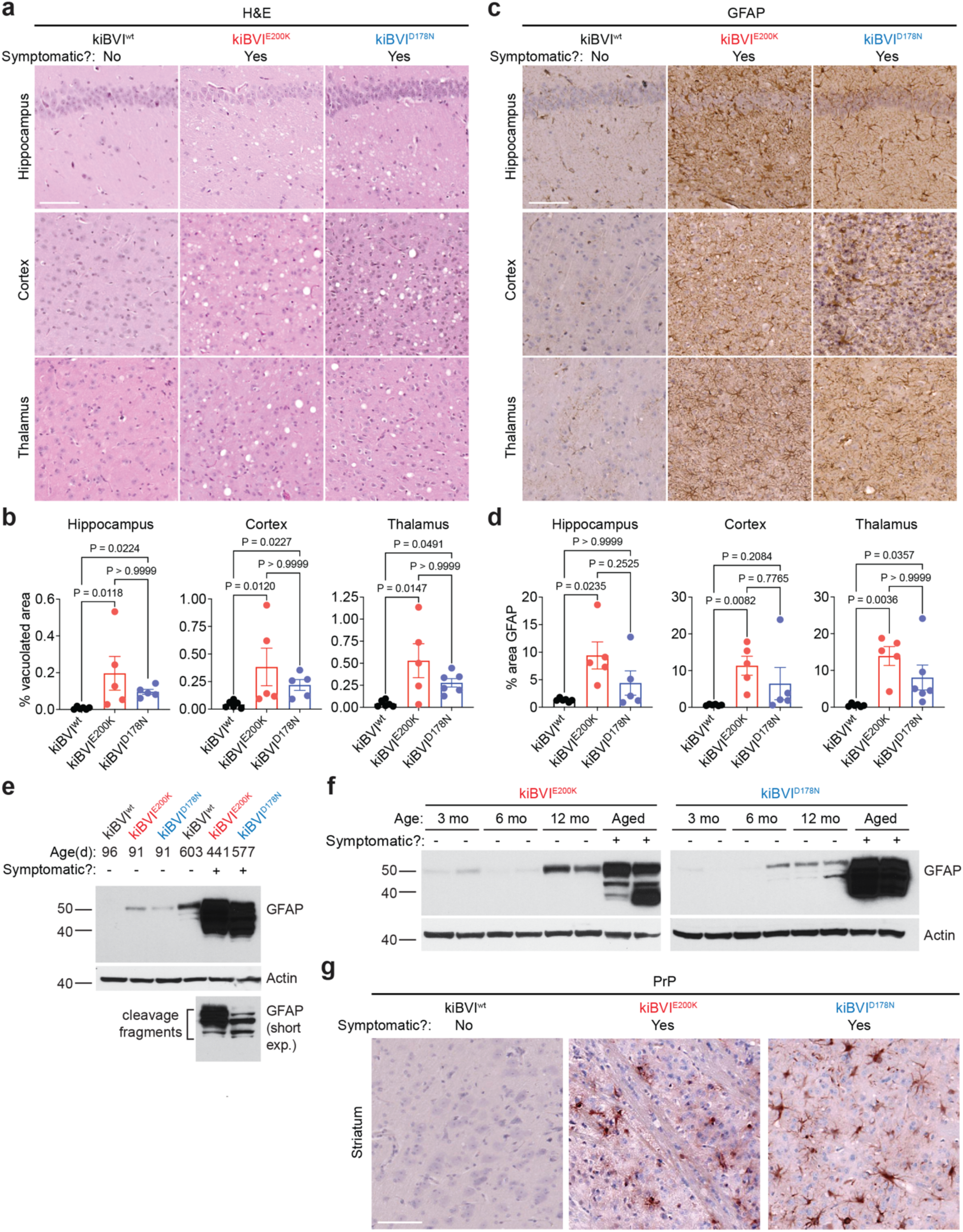
Spontaneously ill knock-in mice expressing mutant bank vole PrP exhibit prion disease-specific neuropathology. **a**) Representative H&E-stained brain sections of the hippocampus, cortex, and thalamus from spontaneously ill kiBVI^E200K^ and kiBVI^D178N^ mice as well as 20-month-old asymptomatic kiBVI^wt^ mice. Scale bar = 100 µm (applies to all sections). **b**) Quantification of percent area covered by vacuolation (mean ± SEM) in the indicated brain regions from aged knock-in mice (n = 5-6 samples per line). **c**) Representative GFAP-stained brain sections of the hippocampus, cortex, and thalamus from spontaneously ill kiBVI^E200K^ and kiBVI^D178N^ mice as well as 20-month-old asymptomatic kiBVI^wt^ mice. Scale bar = 100 µm (applies to all sections). **d**) Quantification of percent area covered by GFAP staining in the indicated brain regions from aged knock-in mice. n = 5-6 samples per line. **e**) Immunoblot of GFAP levels in brain homogenates from the three lines of knock-in mice at the indicated ages. The blot was reprobed with an antibody against actin. **f**) Immunoblots of GFAP levels in brain extracts from kiBVI^E200K^ (left) and kiBVI^D178N^ (right) mice at the indicated ages. Two independent mice per age were analyzed, and the blots were reprobed with an antibody against actin. **g**) Representative PrP-stained brain sections of the striatum from spontaneously ill kiBVI^E200K^ and kiBVI^D178N^ mice as well as 20-month-old asymptomatic kiBVI^wt^ mice. Scale bar = 100 µm (applies to all sections). Molecular weight markers in panels e and f are in kDa. In panels b and d, statistical significance was assessed using a Kruskal-Wallis test followed by Dunn’s multiple comparison test.

Immunohistochemical staining for PrP using the 9A2 antibody failed to detect any extracellular PrP^Sc^ deposition in regions of the brain that exhibited spongiform degeneration from spontaneously ill kiBVI^D178N^ and kiBVI^E200K^ mice. However, intracellular PrP deposition within glial cells was selectively present in the striatum of sick kiBVI^D178N^ and kiBVI^E200K^ mice (**Fig. 4g**), and there were no apparent differences in PrP deposition patterns between the two lines. We hypothesize that this intracellular staining may represent uptake of misfolded PrP species by glial cells and subsequent failure of the endosomal/lysosomal system to clear these aggregates effectively.

### Transmission properties of spontaneously formed prions

To test if prion infectivity was present in the brains of spontaneously ill kiBVI^E200K^ and kiBVI^D178N^ mice, we performed transmission studies in kiBVI^wt^ mice, which do not themselves develop spontaneous disease (**Fig. 5a**). Two inocula each from spontaneously ill kiBVI^E200K^ and kiBVI^D178N^ mice were selected based on the presence of high amounts of TL-resistant PrP in brain extracts. As a negative control, kiBVI^wt^ mice were inoculated with brain extracts prepared from either of two distinct asymptomatic 20-month-old kiBVI^wt^ mice. All the mice inoculated with kiBVI^wt^ extract remained free of neurological illness for up to 18 months post-inoculation (**Fig. 5b**, **Table 2**). To the contrary, 100% of kiBVI^wt^ mice inoculated with brain extracts from spontaneously ill kiBVI^E200K^ mice developed neurological illness with mean incubation periods of ∼10-11 months. The most common clinical signs were kyphosis, weight loss, tremor, and bradykinesia. Disease transmission was less efficient when using inocula prepared from spontaneously ill kiBVI^D178N^ mice, potentially due to a slight transmission barrier introduced by the D178N mutation, with only ∼40% of inoculated mice exhibiting neurological illness within the 18-month experimental timeframe (**Fig. 5b**, **Table 2**).

**Figure 5.**
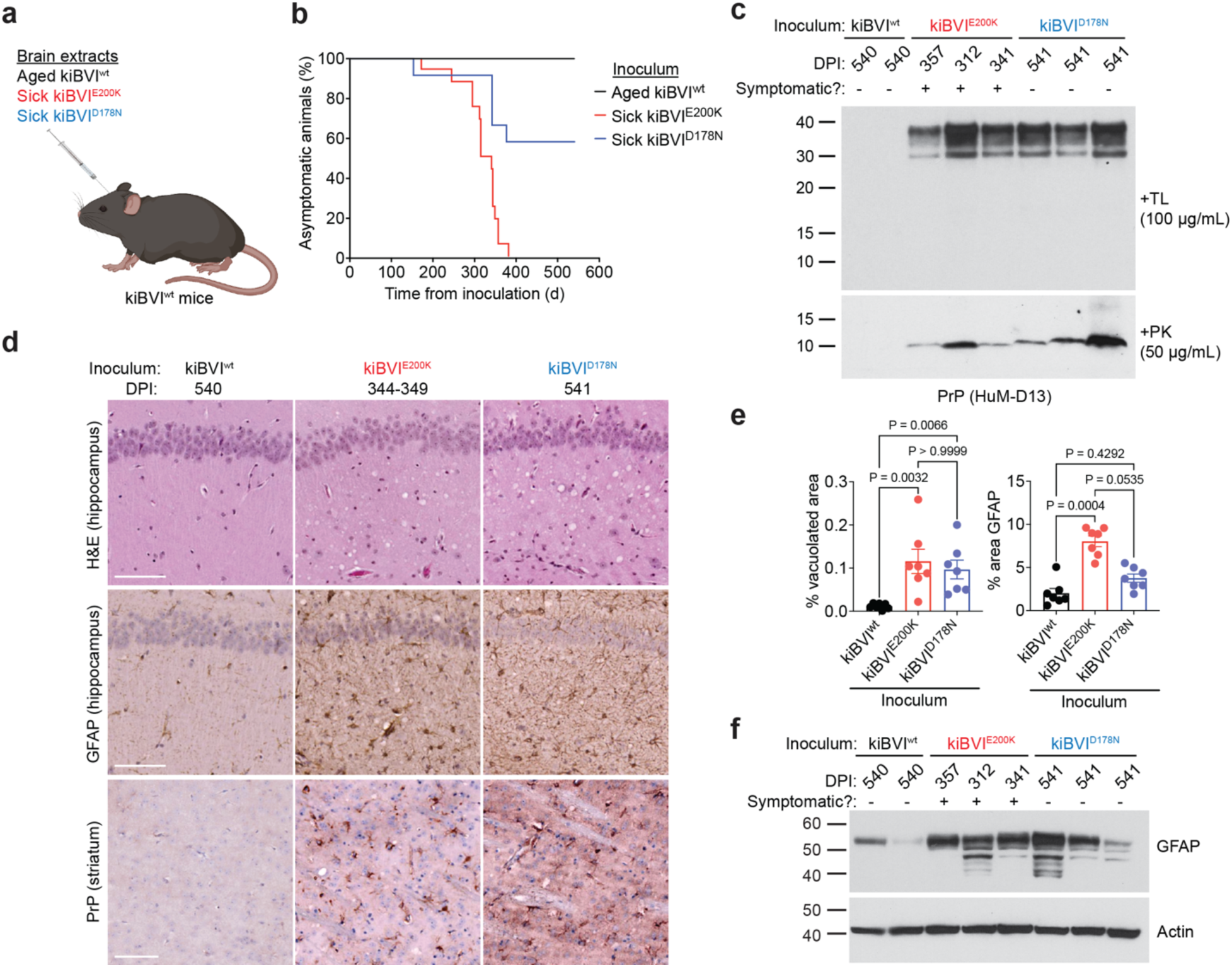
Transmission of prions from the brains of spontaneously ill knock-in mice expressing mutant bank vole PrP to knock-in mice expressing wild-type bank vole PrP. **a**) Schematic of transmission experiments in kiBVI^wt^ mice. **b**) Kaplan-Meier curves for the development of signs of neurological illness in kiBVI^wt^ mice inoculated with brain extract from aged, asymptomatic kiBVI^wt^ mice (black, n = 17), symptomatic kiBVI^E200K^ mice (red, n = 16), or symptomatic kiBVI^D178N^ mice (blue, n = 13). The onset of neurological illness in mice inoculated with kiBVI^E200K^ or kiBVI^D178N^ extract was significantly different from mice inoculated with kiBVI^wt^ extract (*P* < 0.0001 and *P* = 0.0036, respectively, as assessed by the Log-rank test). **c**) Immunoblots for detergent-insoluble PrP species in brain extracts from kiBVI^wt^ mice at the indicated days post-inoculation (DPI) with brain extract from kiBVI^wt^ mice, symptomatic kiBVI^E200K^ mice, or symptomatic kiBVI^D178N^ mice following digestion with either 100 µg/mL TL (top panel) or 50 µg/mL PK (bottom panel). TL- and PK-resistant PrP species were detected using the antibody HuM-D13. **d**) Representative H&E- and GFAP-stained sections of the hippocampus and PrP-stained sections of the striatum from kiBVI^wt^ mice at the indicated DPI with brain extract from either spontaneously ill kiBVI^E200K^ or kiBVI^D178N^ mice or 20-month-old asymptomatic kiBVI^wt^ mice. Scale bars = 100 µm. **e**) Quantification of percent area covered by vacuolation or GFAP staining in the inoculated kiBVI^wt^ mice. n = 7 samples per line. Statistical significance was assessed using a Kruskal-Wallis test followed by Dunn’s multiple comparison test. **f**) Immunoblot of GFAP levels in brain extracts from kiBVI^wt^ mice at the indicated DPI with brain extract from kiBVI^wt^ mice, symptomatic kiBVI^E200K^ mice, or symptomatic kiBVI^D178N^ mice. The blot was reprobed with an antibody against actin. Molecular weight markers in panels c and f are in kDa.

**Table 2.**
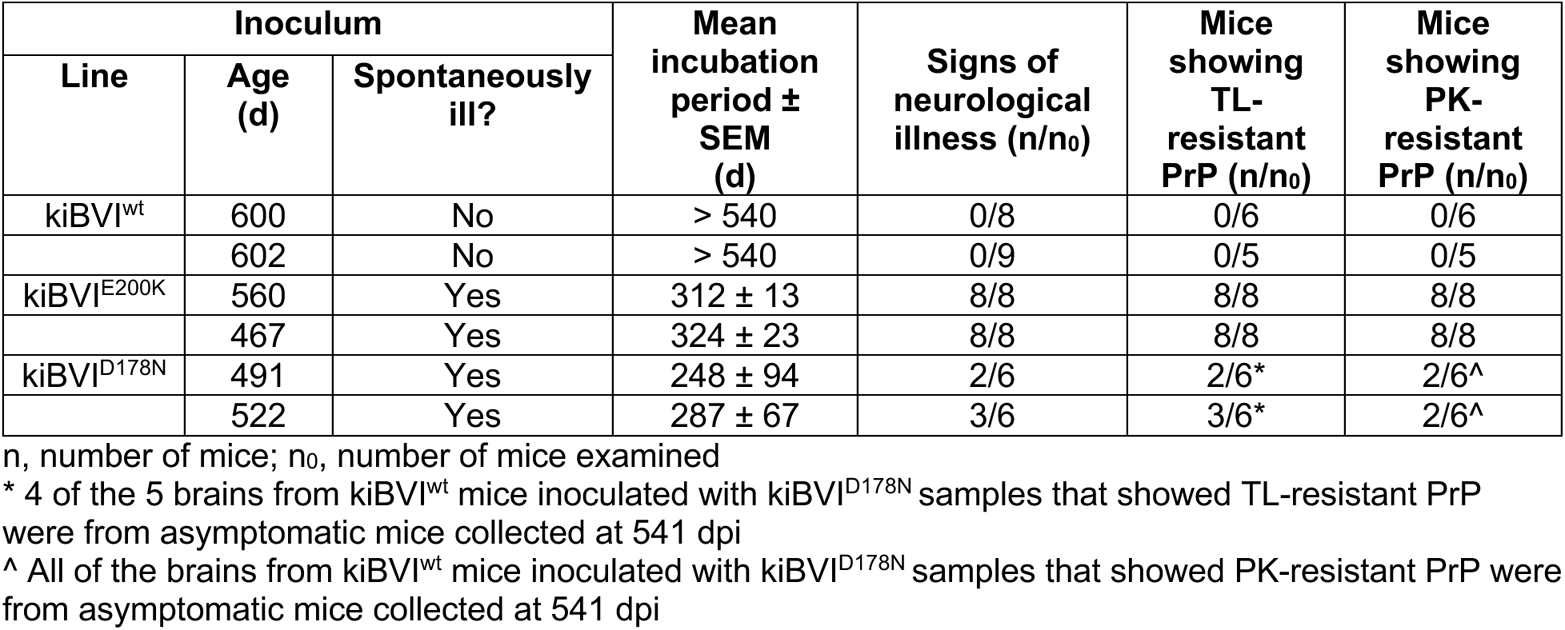
Transmission of brain homogenates from BVPrP knock-in mice to kiBVI^wt^ mice.

| Inoculum | | | Mean incubation period ± SEM (d) | Signs of neurological illness (n/n<sub>0</sub>) | Mice showing TL-resistant PrP (n/n<sub>0</sub>) | Mice showing PK-resistant PrP (n/n<sub>0</sub>) |
| --- | --- | --- | --- | --- | --- | --- |
| Line | Age (d) | Spontaneously ill? |  |  |  |  |
| kiBVI<sup>wt</sup> | 600 | No | > 540 | 0/8 | 0/6 | 0/6 |
|  | 602 | No | > 540 | 0/9 | 0/5 | 0/5 |
| kiBVI<sup>E200K</sup> | 560 | Yes | 312 ± 13 | 8/8 | 8/8 | 8/8 |
| | 467 | Yes | 324 ± 23 | 8/8 | 8/8 | 8/8 |
| kiBVI<sup>D178N</sup> | 491 | Yes | 248 ± 94 | 2/6 | 2/6* | 2/6^ |
| | 522 | Yes | 287 ± 67 | 3/6 | 3/6* | 2/6^ |

n, number of mice; n<sub>0</sub>, number of mice examined
\* 4 of the 5 brains from kiBVI<sup>wt</sup> mice inoculated with kiBVI<sup>D178N</sup> samples that showed TL-resistant PrP were from asymptomatic mice collected at 541 dpi
^ All of the brains from kiBVI<sup>wt</sup> mice inoculated with kiBVI<sup>D178N</sup> samples that showed PK-resistant PrP were from asymptomatic mice collected at 541 dpi

Whereas brains from kiBVI^wt^ mice inoculated with kiBVI^wt^ extract did not contain PrP^res^, both TL- and PK-resistant PrP species were detected in the brains of all mice inoculated with kiBVI^E200K^ extract (**Fig. 5c**). Likewise, TL- and PK-resistant PrP were found in the brains of some, but not all kiBVI^wt^ mice inoculated with kiBVI^D178N^ extract. Surprisingly, of the 5 kiBVI^wt^ brains that displayed TL-resistant PrP following inoculation with kiBVI^D178N^ extract, 4 were from mice collected at 540 days post-inoculation in the absence of clinical signs of neurological illness. The TL- and PK-resistant PrP species in the inoculated kiBVI^wt^ mice were reminiscent of those present in spontaneously ill kiBVI^E200K^ and kiBVI^D178N^ mice, and there were no obvious differences in molecular weights of the PrP^res^ fragments induced by the kiBVI^E200K^ and kiBVI^D178N^ lines. The brains of kiBVI^wt^ mice inoculated with kiBVI^E200K^ or kiBVI^D178N^ extract also contained neuropathological indicators of prion disease such as vacuolation and astrocytic gliosis as well as increased endoproteolytic cleavage of GFAP (**Fig. 5d-f**). Unlike in the spontaneously sick mice, vacuolation in the kiBVI^wt^ mice inoculated with kiBVI^E200K^ or kiBVI^D178N^ samples was largely restricted to the hippocampus. GFAP staining in the hippocampus was less pronounced in the mice inoculated with samples from kiBVI^D178N^ mice, likely reflecting the reduced rate of symptomatic transmission. Intracellular PrP deposition within striatal glial cells of inoculated kiBVI^wt^ mice was observed in animals inoculated with brain extract from kiBVI^E200K^ or kiBVI^D178N^ mice (**Fig. 5d**). There were no obvious differences in the neuropathological signatures of kiBVI^wt^ mice inoculated with kiBVI^E200K^ or kiBVI^D178N^ samples. Collectively, these results demonstrate that transmissible prions can form spontaneously in the brains of kiBVI^E200K^ and kiBVI^D178N^ mice, but not in the brains of kiBVI^wt^ mice.

### Conformational characterization of spontaneous and transmitted prions

To characterize the conformational properties of the prions formed spontaneously in the brains of kiBVI^E200K^ and kiBVI^D178N^ mice, we employed a conformational stability assay that measures the relative resistance of protein aggregates to denaturation with guanidine hydrochloride (**Fig. S6**) [45]. This assay is commonly used to discriminate between different prion strains [46, 65]. There was no difference in the conformational stability of the TL-resistant PrP aggregates in the brains of spontaneously ill kiBVI^E200K^ and kiBVI^D178N^ mice (**Fig. 6a, b**). Similarly, no differences in conformational stability were observed following transmission to kiBVI^wt^ mice (**Fig. 6c, d**).

**Figure 6.**
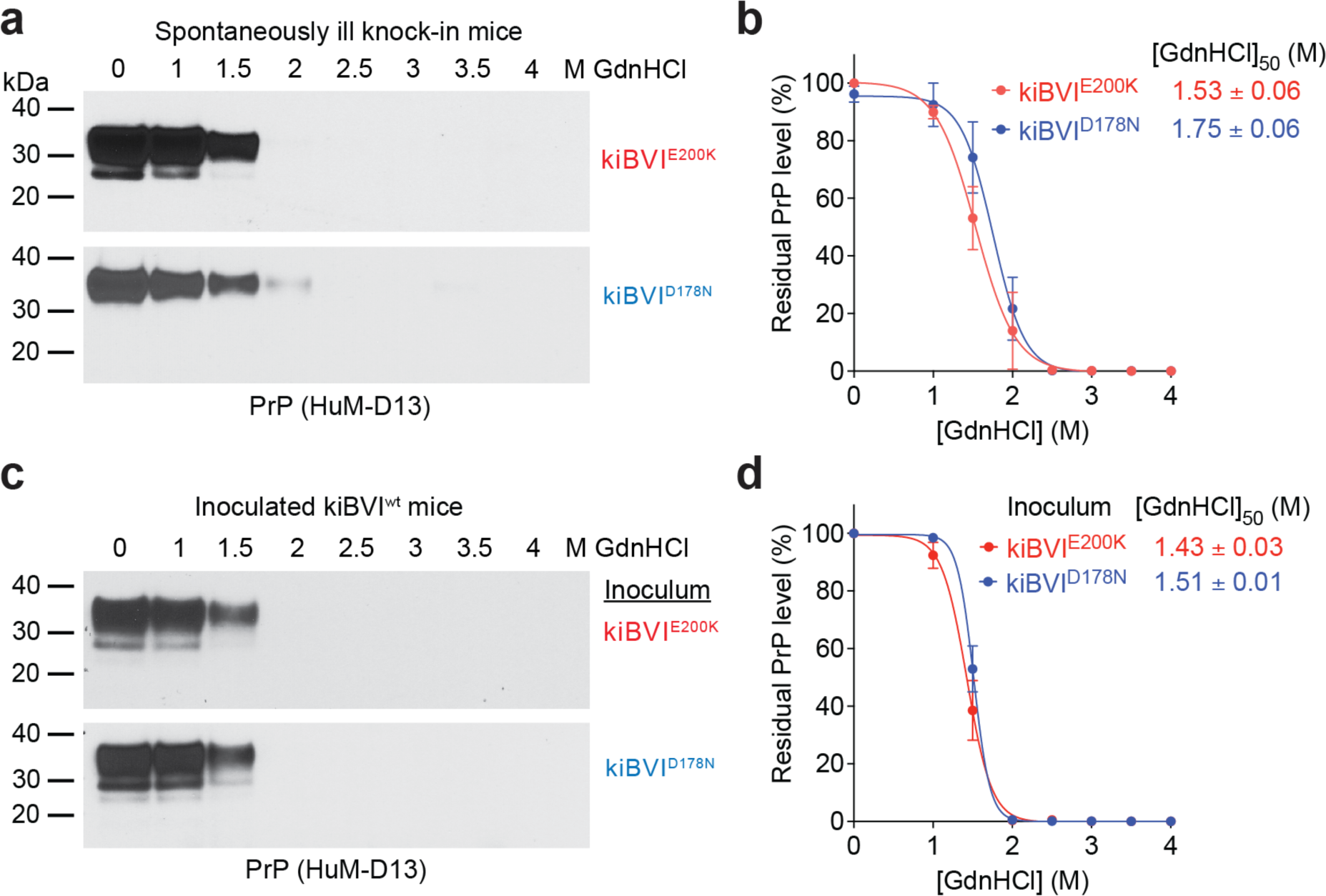
Conformational characterization of spontaneously formed and transmitted prions. **a**) Representative immunoblots of detergent-insoluble, TL-resistant PrP species following treatment of brain extracts from spontaneously ill kiBVI^E200K^ mice (top blot) or kiBVI^D178N^ mice (bottom blot) with the indicated concentrations of guanidine hydrochloride (GdnHCl). **b**) Quantification of residual detergent-insoluble, TL-resistant PrP levels in brain extracts from spontaneously ill kiBVI^E200K^ mice (red, n = 3) or kiBVI^D178N^ mice (blue, n = 3) treated with the indicated concentrations of GdnHCl. **c**) Representative immunoblots of detergent-insoluble, TL-resistant PrP species in brain homogenates from kiBVI^wt^ mice inoculated with either kiBVI^E200K^ (top blot) or kiBVI^D178N^ (bottom blot) brain extract treated with the indicated concentrations of GdnHCl. **d**) Quantification of residual detergent-insoluble, TL-resistant PrP levels in brain homogenates from kiBVI^wt^ mice inoculated with brain extract from either kiBVI^E200K^ (red, n = 3) or kiBVI^D178N^ (blue, n = 3) mice following treatment with the indicated concentrations of GdnHCl. In panels a and c, PrP was detected using the antibody HuM-D13. In panels b and d, the calculated [GdnHCl]_50_ values are also shown.

## Discussion

To the best of our knowledge, the kiBVI^D178N^ and kiBVI^E200K^ mice described herein are the first mouse models that exhibit all the cardinal features of human prion diseases without requiring the use of PrP overexpression, a non-native promoter, or injection with a pre-existing source of PrP^Sc^. The mice developed progressive signs of neurological illness, prion disease-specific neuropathological changes such as spongiform degeneration and astrocytic gliosis, as well as TL- and PK-resistant PrP species in their brains. Most importantly, brain extracts from spontaneously sick mice expressing mutant BVPrP transmitted disease to kiBVI^wt^ mice expressing wild-type BVPrP, which did not develop spontaneous disease themselves, confirming the generation of authentic prion infectivity in the kiBVI^D178N^ and kiBVI^E200K^ lines. Thus, these mice will be useful for interrogating the biological mechanisms governing spontaneous prion formation in the brain during FFI and fCJD as well as uncovering therapeutic strategies for counteracting these processes. For instance, we show in a complementary study that small molecules known to interfere with templated prion replication do not prolong the onset of spontaneous disease in either the kiBVI^D178N^ or kiBVI^E200K^ lines [83]. A limitation of our study is that while the D178N and E200K mutations are most commonly heterozygous in cases of genetic prion disease (i.e., wt and mutant PrP are simultaneously present), we employed mice that were homozygous for the mutations and thus expressed higher relative levels of mutant PrP. However, it should be noted that individuals with homozygous E200K mutations have been identified and develop typical fCJD but at a younger age than those with a heterozygous mutation [57, 76].

We utilized the I109 polymorphic variant of BVPrP for our studies because it more readily forms prions spontaneously when overexpressed in transgenic mice than the M109 polymorphic variant [86]. Indeed, transgenic mice that display modest overexpression of sheep PrP with isoleucine at position 112, which is analogous to position 109 in BVPrP, also develop a spontaneous and transmissible prion disease [82]. Transgenic mice expressing very high levels of the M109 BVPrP polymorphic variant developed a late-onset, but non-transmissible spontaneous neurological illness characterized by spongiform degeneration and PrP deposition in the absence of PrP^res^ [41]. Overexpression of mouse or hamster PrP in transgenic mice can also lead to a proteinopathy that resembles authentic prion disease but is non-transmissible [37, 89]. Thus, the presence of isoleucine at codon 109 of BVPrP may be critical for permitting the formation of PrP assemblies that are able to self-propagate. However, we did not observe any signs of spontaneous prion formation in aged kiBVI^wt^ mice, which is consistent with the lack of reports describing spontaneous disease in I109 bank voles. Therefore, PrP overexpression is required to initiate spontaneous prion formation in mice expressing wt PrP, even when a permissive substrate such as BVPrP is utilized.

Knock-in mouse models that express D178N- or E200K-mutant MoPrP have previously been generated and spontaneously develop prions that can be transmitted to mice expressing wild-type MoPrP [38, 39]. However, unlike the BVPrP-based knock-ins, they do not develop overt signs of progressive neurological illness and fail to exhibit the PrP^res^ species that typify authentic prion disease. Besides the presence of isoleucine at codon 109, the sequence determinants of BVPrP that allow it to function as a superior substrate for studying spontaneous prion formation compared to MoPrP remain unknown. One study found that the addition of two BVPrP-specific C-terminal residues (E227 and S230) to MoPrP was sufficient to elicit spontaneous disease in mice [41], and this region has been implicated as a “linchpin domain” responsible for BVPrP’s enhanced susceptibility to prion strains from other species [12]. Other studies have identified residues within the α-helical domain of BVPrP, specifically asparagine at positions 155 and 170, that may facilitate cross-species prion transmission [1, 43]. How any of these specific residue differences may promote prion formation is unclear, since the structure of bank vole PrP^C^ is very similar to that of PrP^C^ from other mammals, other than the existence of a more rigid loop immediately prior to the second α-helix [16]. The presence of a rigid loop in the structure of PrP^C^ has been linked to the appearance of spontaneous prion generation in transgenic mice [74, 75].

While brains from most aged knock-in mice expressing mutant BVPrP exhibited TL-resistant PrP, a smaller fraction contained PK-resistant PrP. This suggests that a spectrum of PrP assemblies develop in the knock-in mice, with the formation of TL-resistant species likely preceding the generation of potentially larger or more densely packed PK-resistant aggregates (**Fig. 7**). Whereas FFI and fCJD patients develop stereotypical PrP^res^ with a molecular weight of 19-21 kDa following removal of N-linked glycans [55, 80], the brains of spontaneously sick kiBVI^D178N^ and kiBVI^E200K^ mice exhibited a smaller PK-resistant PrP fragment with a molecular weight of ∼10 kDa, which is more reminiscent of those found in GSS [66, 77, 78]. Short PK-resistant PrP fragments can also be found in Nor98/atypical scrapie in sheep and in variably protease-sensitive prionopathy (VPSPr) in humans [8, 69, 93]. Interestingly, these “atypical” prion diseases, all of which are thought to result from spontaneous misfolding of PrP, can be efficiently transmitted to bank voles with the I109 polymorphism [59, 67, 68], and ∼7 kDa non-fibrillar PrP^res^ species purified from GSS brains are sufficient for disease transmission [81].Thus, the presence of the I109 residue may selectively stabilize these misfolded PrP species, allowing them to propagate and spread within the brain. The presence of prion disease-associated neuropathological changes such as spongiform degeneration in spontaneously ill kiBVI^D178N^ and kiBVI^E200K^ mice despite the absence of extracellular PrP^Sc^ deposition and stereotypical PrP^res^ is consistent with the hypothesis that stereotypical PrP^res^ may not be the principal neurotoxic species in prion disease [9, 72].

**Figure 7.**
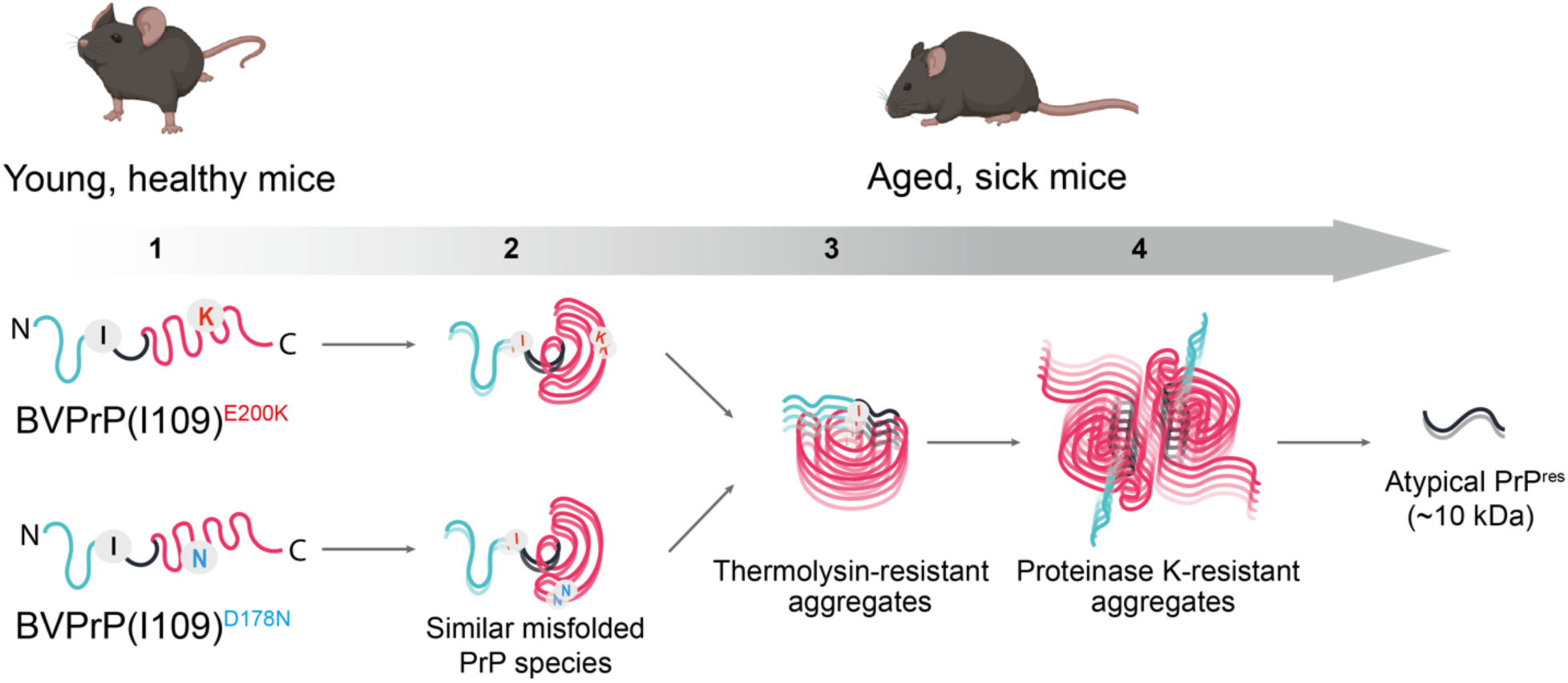
Model for the spontaneous generation of prions in kiBVI^E200K^ and kiBVI^D178N^ mice. (1) Properly folded PrP is present in the brains of young, healthy knock-in mice expressing mutant BVPrP(I109). (2) As the mice age, the E200K and D178N mutations promote the generation of a very similar misfolded BVPrP species. (3) Polymerization of this misfolded species results in BVPrP aggregates that are resistant to thermolysin digestion and the appearance of clinical signs of neurological illness in the knock-in mice. (4) As disease progresses, the BVPrP aggregates undergo conformational maturation to produce aggregates that contain a core region that is resistant to proteinase K digestion, producing an atypical PrP^res^ fragment of ∼10 kDa. This PrP^res^ fragment does not contain either the D178N- or E200K-mutant residues.

Potential relationships between atypical PrP^res^ and the stereotypical PrP^res^ species found in most animal and human prion diseases remain to be fully investigated. Repeated passage of Nor98/atypical scrapie in transgenic mice expressing bovine PrP leads to a molecular phenotype indistinguishable from classical bovine spongiform encephalopathy, which is typified by the presence of stereotypical PrP^res^ [36]. Moreover, whereas I109 bank voles faithfully propagate the atypical short PrP^res^ fragments upon transmission of Nor98 scrapie, Nor98-inoculated M109 bank voles exhibit classical PrP^res^ [68]. Intriguingly, cases of GSS caused by the P102L mutation exhibit a short ∼8 kDa PrP^res^ fragment, either alone or in combination with stereotypical PrP^res^ [54, 63]. Therefore, stereotypical PrP^res^ may emerge from or along with atypical PrP^res^, potentially due to selection of rare conformational species within a mixture or deformed templating [18, 47].

The striking similarities in the neuropathological profiles, PrP^res^ signatures, conformational stabilities, and transmission properties of the prions present in kiBVI^D178N^ and kiBVI^E200K^ mice imply that both lines spontaneously develop a highly similar, if not identical atypical prion strain. This was unexpected given that BVPrP-based transgenic models of FFI and fCJD exhibit mutation-specific pathological signatures, although this could potentially be an artifact of PrP over-expression or differences in spatial expression patterns [84]. MoPrP-based knock-in models of FFI and fCJD also exhibit different pathological phenotypes, perhaps due to the lack of residue I109 since MoPrP contains leucine at corresponding residue 108 [38, 39]. However, translatome studies have shown that the molecular phenotypes in the MoPrP-based FFI and fCJD knock-in lines are more similar than expected [7]. Although we cannot rule out a scenario where the properties of BVPrP(I109) override mutation-specific strain generation, our data raise the possibility that the D178N and E200K mutations may uniformly act to promote the initial misfolding of PrP into a similar self-propagating species (**Fig. 7**). The key properties of this initial misfolded species are likely determined by its atypical PK-resistant core rather than by the specific mutation, since similar fragments are known to be both N- and C-terminally truncated and do not contain either residue 178 or 200 [30, 69, 81]. The mechanism by which the D178N and E200K mutations promote the accumulation of misfolded BVPrP remains to be established. The mutations may either directly promote spontaneous misfolding of BVPrP, stabilize the formation of prion assemblies, and/or reduce the clearance rates of aggregates. An increased turnover rate for the mutant proteins implies a mutation-induced decrease in protein stability [84], and the D178N mutation may further accelerate misfolding by destabilization of the C1 endoproteolytic fragment, which is thought to be an endogenous inhibitor of prion conversion [90].

In humans, disease-causing mutations within PrP clearly influence prion strain generation in the brain. For instance, individuals with the D178N mutation coupled to M129 only develop FFI. In contrast, those with the E200K-M129 haplotype develop CJD with one of three distinct PrP^res^ signatures [6]. We propose a dual-hit model in which PrP mutations play two roles in genetic prion disease. First, they act by promoting the accumulation of an atypical misfolded PrP species and second, later in the disease course, they bias the emergence of classical PrP^res^ towards conformations that are compatible with the specific mutation. Given that this latter phase might require a rare secondary misfolding event, our two-phase model can provide insights into why genetic prion diseases manifest later in life, even though the mutations exist from birth. Facilitated by the inherent misfolding propensity of BVPrP and the capacity of I109 to stabilize atypical prion species, kiBVI^D178N^ and kiBVI^E200K^ mice may effectively recapitulate pivotal early PrP misfolding events that occur in the brain during genetic prion disease.

## Acknowledgements

We are grateful to Sue Plyte and the staff at the UHN Animal Resources Centre for their assistance with facilitating the prion inoculation experiments. The authors thank Dr. Stanley Prusiner (University of California San Francisco) for providing the HuM-D13 antibody and the HRP-labeled HuM-P antibody. This work was funded by a grant from the Canadian Institutes of Health Research (PJT-169048). SM was supported by a postdoctoral fellowship from Parkinson Canada.

## Declarations

### Ethics approval

All mouse experiments were performed in accordance with guidelines set by the Canadian Council on Animal Care under protocols (AUP #4263.17 and 6322.3) approved by the University Health Network Animal Care Committee.

### Availability of data and material

All data generated or analyzed during this study are included in this published article.

### Author’s contributions

All authors contributed to the study conception and design. Material preparation, data collection and analysis were performed by Surabhi Mehra, Matthew E.C. Bourkas, Lech Kaczmarczyk, Erica Stuart, Hamza Arshad, Jennifer K. Griffin, Kathy L. Frost, Daniel J. Walsh, Surachai Supattapone, Stephanie A. Booth, Walker S. Jackson, and Joel C. Watts. The first draft of the manuscript was written by Surabhi Mehra, Matthew E.C. Bourkas and Joel C. Watts, and all authors commented on previous versions of the manuscript. All authors read and approved the final manuscript.

