## Supplemental data for "Convergent generation of atypical prions in knock-in mouse models of genetic prion disease"

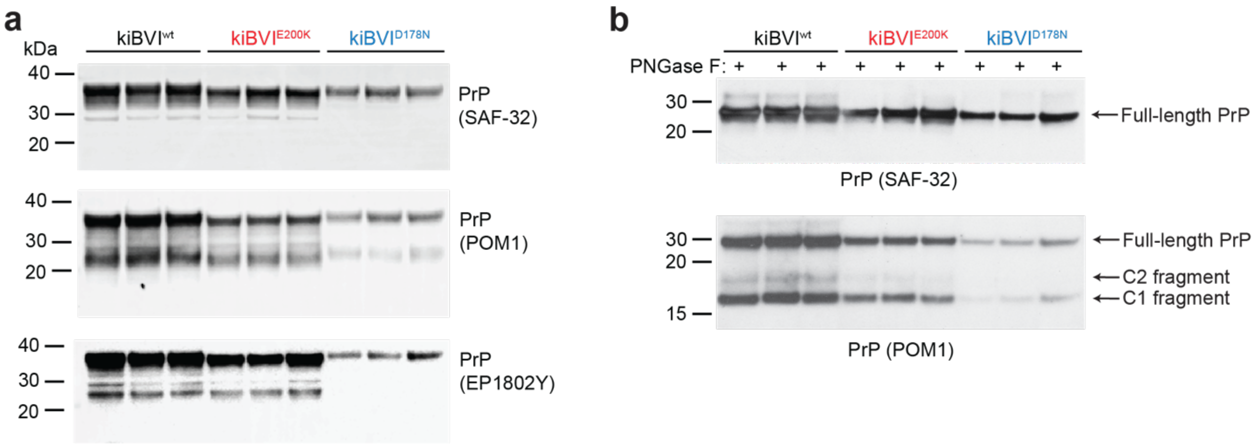


**Fig. S1** Characterization of knock-in mice expressing wild-type or mutant bank vole PrP using additional antibodies. **a**) Immunoblots for PrP in brain extracts from 3 mice each for the kiBVI$^{wt}$, kiBVI$^{E200K}$, and kiBVI$^{D178N}$ lines. BVPrP was detected using the antibodies SAF-32, POM1, and EP1802Y. **b**) Immunoblots for PrP in PNGase F-treated brain extracts from 3 mice each for the kiBVI$^{wt}$, kiBVI$^{E200K}$, and kiBVI$^{D178N}$ lines. BVPrP was detected using the antibodies SAF-32 and POM1. Full-length BVPrP as well as the C1 and C2 endoproteolytic products are indicated

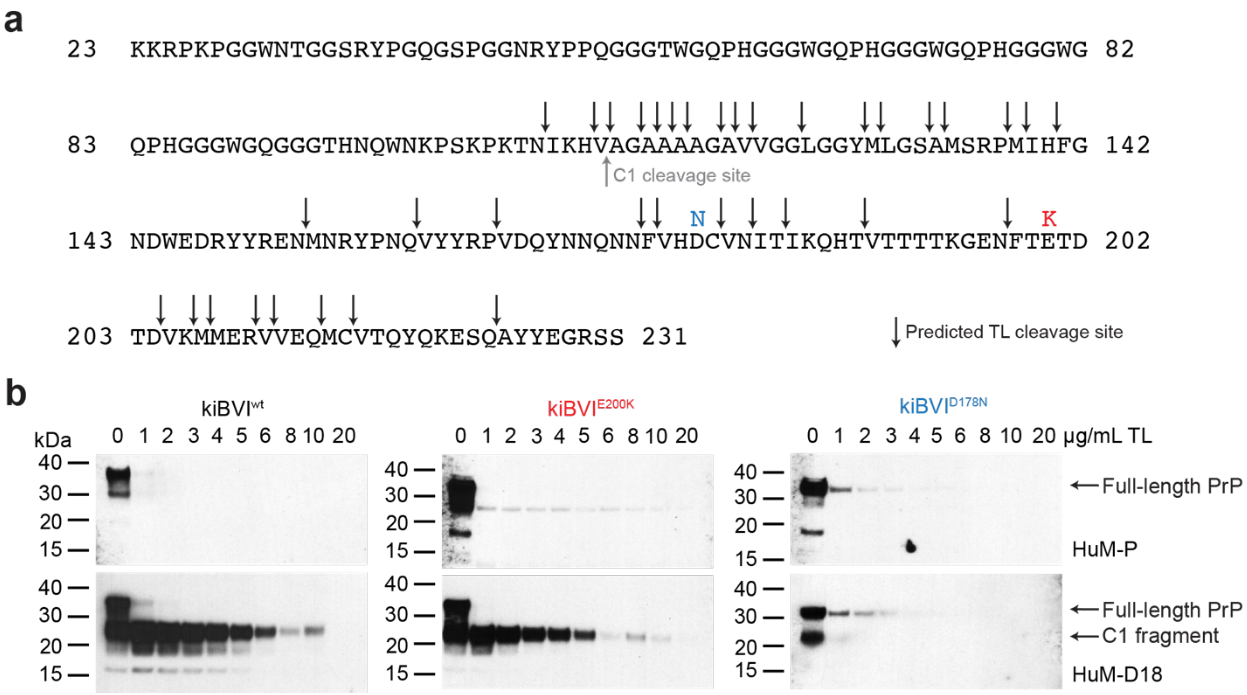


**Fig. S2** Thermolysin digestion of brain extracts from young, asymptomatic knock-in mice. **a**) Predicted thermolysin (TL) cleavage sites in the sequence of the mature form of BVPrP lacking the N- and C-terminal signal sequences. The locations of the D178N and E200K mutations as well as the C1 endoproteolytic cleavage site are also shown. **b**) Immunoblots for PrP in brain extracts from young (3-month-old) kiBVI$^{wt}$ (left), kiBVI$^{E200K}$ (middle), and kiBVI$^{D178N}$ (right) mice treated with the indicated concentrations of TL. Blots were probed with the antibodies HuM-P (top blots) or HuM-D18 (bottom blots). The location of full-length and C1 PrP species are indicated

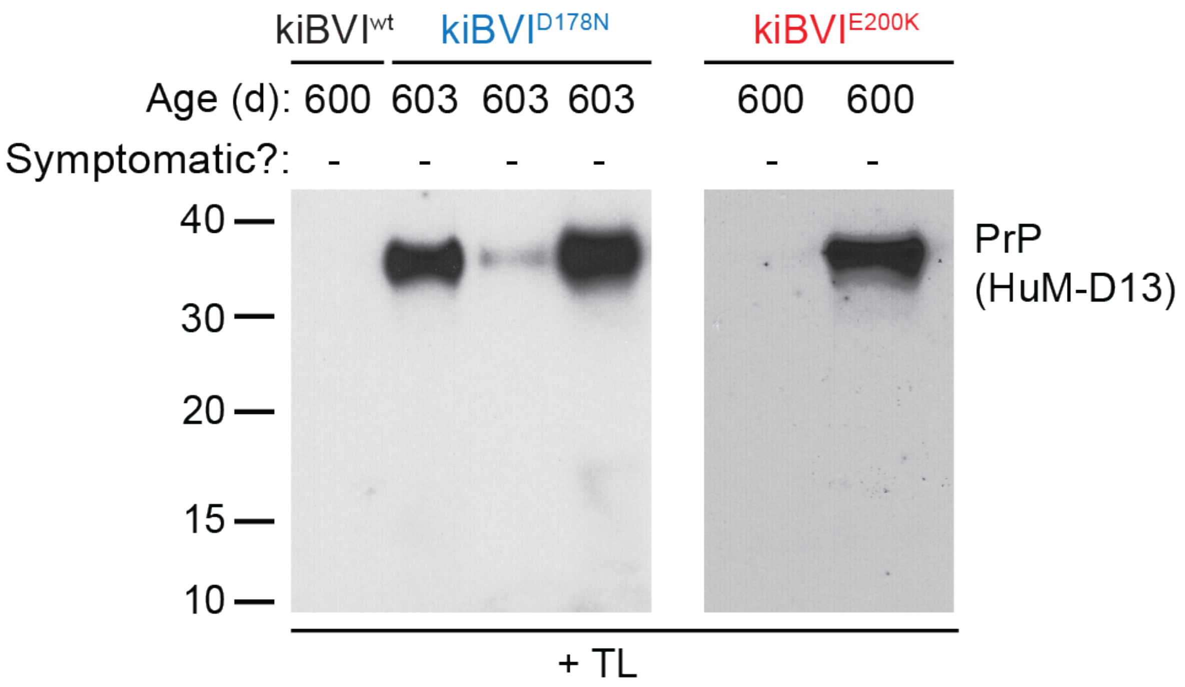


**Fig. S3** Thermolysin-resistant PrP in aged, asymptomatic mice expressing mutant bank vole PrP**.** Immunoblots for detergent-insoluble, TL-resistant PrP species in brain extracts from asymptomatic, 20-month-old kiBVI$^{wt}$, kiBVI$^{D178N}$, and kiBVI$^{E200K}$ mice. PrP was detected using the antibody HuM-D13

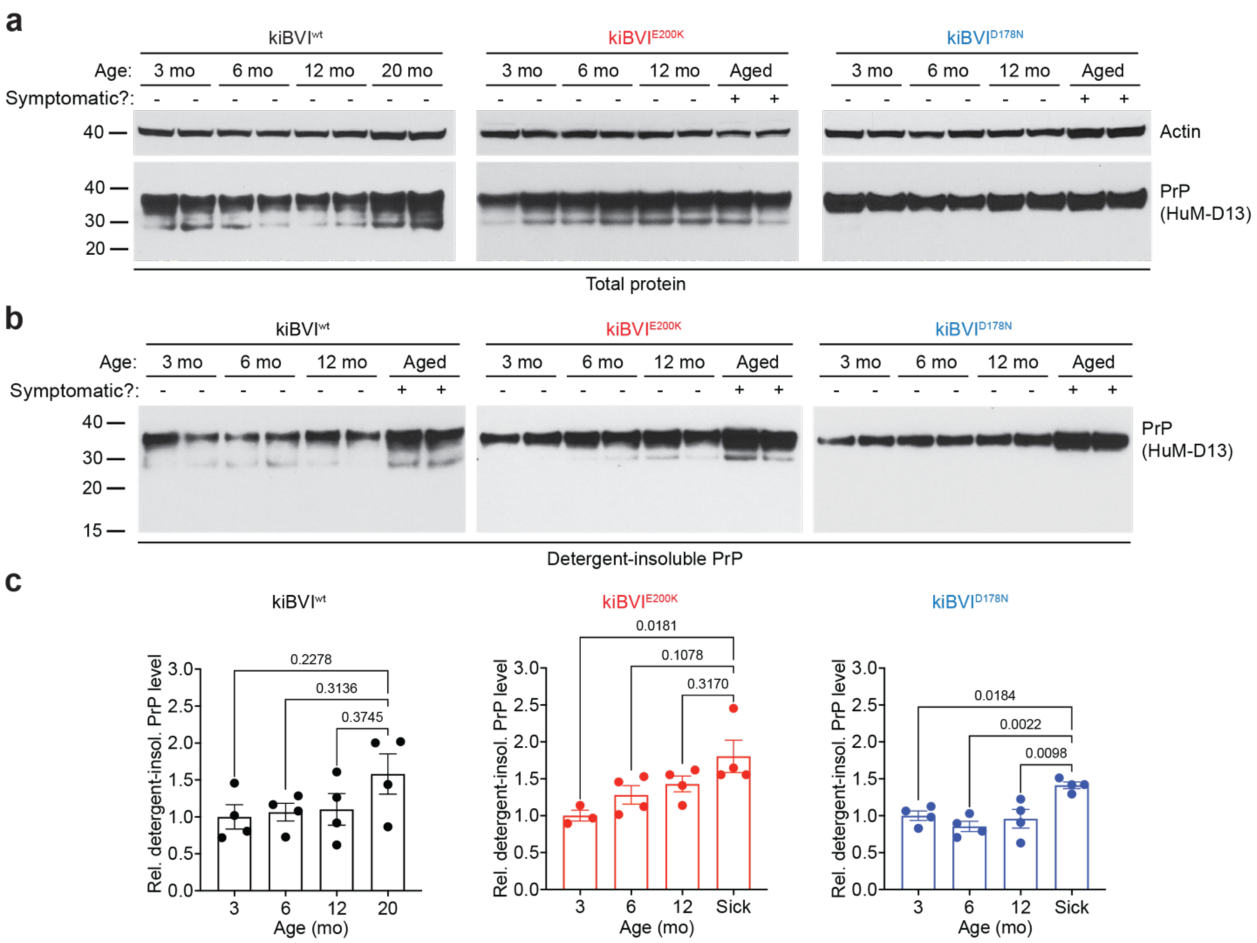


**Fig. S4** Detergent-insoluble PrP levels in knock-in mice at different ages. **a**) Immunoblots of total PrP levels in brain extracts from kiBVI$^{wt}$ (left), kiBVI$^{E200K}$ (middle), and kiBVI$^{D178N}$ (right) mice at the indicated ages. Two independent mice per age were analyzed. For the kiBVI$^{E200K}$ and kiBVI$^{D178N}$ lines, aged mice with spontaneous neurological illness were also examined. **b**) Immunoblots for detergent-insoluble PrP species in brain extracts from kiBVI$^{wt}$ (left), kiBVI$^{E200K}$ (middle), and kiBVI$^{D178N}$ (right) mice at the indicated ages. Two independent mice per age were analyzed. PrP was detected using the antibody HuM-D13. **c**) Quantification of detergent-insoluble PrP species in the three lines at various ages relative to the levels present in the brains of 3-month-old mice for each line (n = 3-4 mice per age). Statistical significance was assessed using one-way ANOVA followed by Tukey's multiple comparisons test

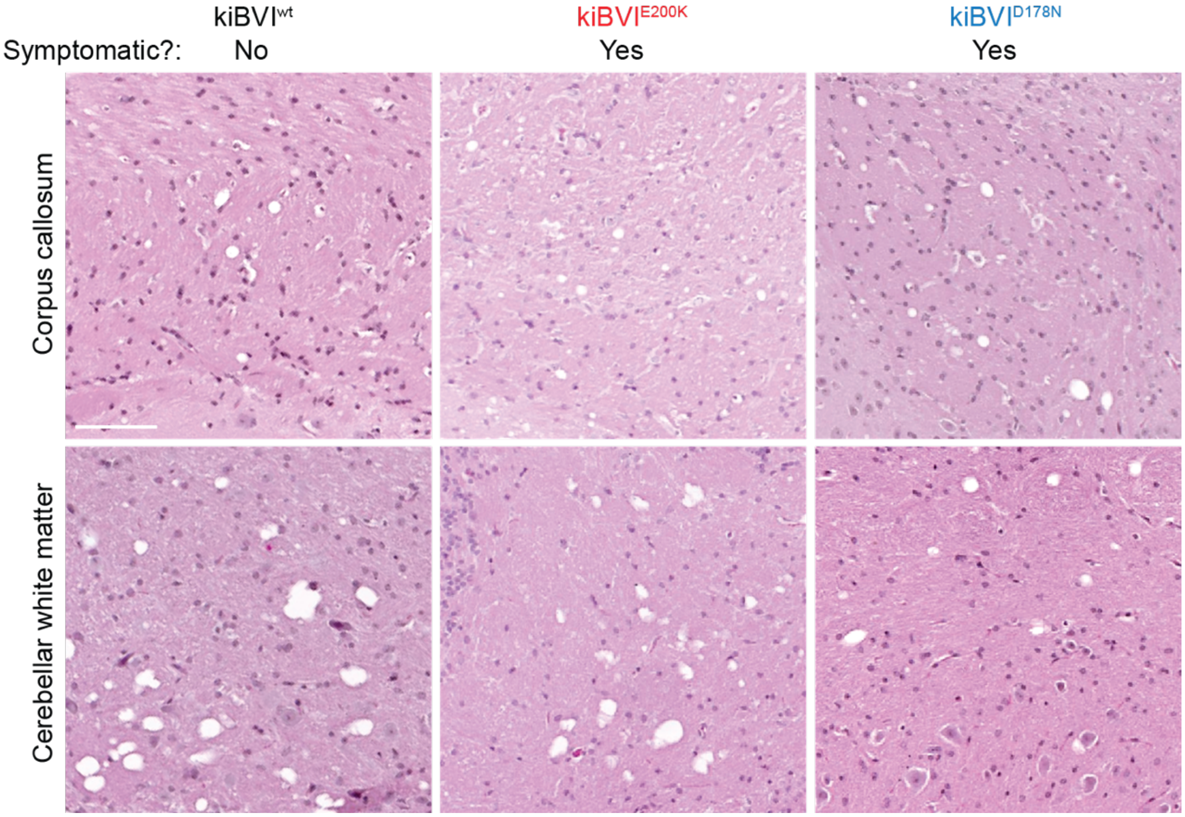


**Fig. S5** White matter vacuolation in aged knock-in mice expressing wild-type or mutant bank vole PrP. Representative H&E-stained sections of the corpus callosum and cerebellar white matter from 20-month-old asymptomatic kiBVI$^{wt}$ mice as well as spontaneously ill kiBVI$^{E200K}$ and kiBVI$^{D178N}$ mice. Scale bar = 100 µm (applies to all sections)

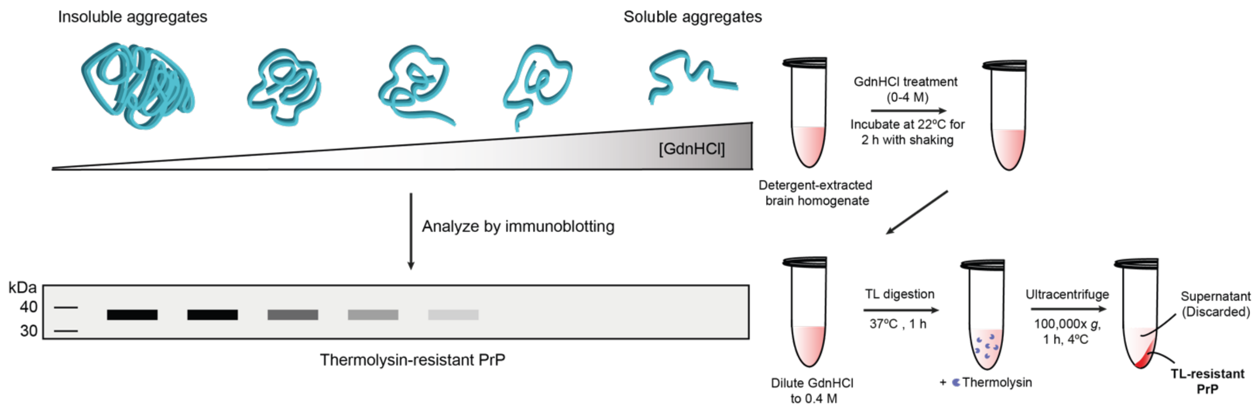


**Fig. S6** Experimental schematic of the conformational stability assay. Following treatment with various concentrations of GdnHCl, samples were digested with thermolysin and then the relative level of insoluble BVPrP species was quantified by immunoblotting
